# Dietary sugar type determines the response to protein restriction in females, but not males

**DOI:** 10.64898/2026.08.18.745207

**Authors:** Michelle M. Sonsalla, Mari Cole, Madeline Johnson, Samuel Cai, Brenna Virnig, Alexandra Trebil, Reji Babygirija, Julia Illiano, Diana Vertein, Yang Liu, Isaac Grunow, Bailey A. Knopf, Sophia Schlorf, Michael Rigby, Chung-Yang Yeh, Cara L. Green, David A. Harris, Luigi Puglielli, Dudley W. Lamming

## Abstract

Low protein (LP) diets improve metabolic health in rodents and humans. In rodents, LP diets are typically implemented by replacing protein with carbohydrates like sucrose or cornstarch, keeping diets isocaloric. However, humans can choose from many different types of carbohydrate, and how dietary carbohydrate quality – the precise composition of the dietary sugars – impacts the response to dietary protein remains largely unexplored. Here, mice were fed control (21% protein) or LP (7% protein) diets with four different carbohydrate sources: sucrose, a 1:1 glucose/fructose mixture, glucose, or fructose. While LP diets improved metabolic health across all groups in male mice, carbohydrate quality also significantly altered specific health outcomes, with fructose-fed mice having the lowest body weight and adiposity of all control diets. In female mice, responses to LP diets were influenced by carbohydrate quality, with certain sugars inducing a stronger metabolic response to LP diets than previously seen. Finally, in female APP/PS1 mice, a model of Alzheimer’s disease, we find that although LP diets reduce Aβ plaque burden irrespective of carbohydrate type, dietary sugar type does influence spatial memory. Together, these results demonstrate that while dietary protein is a critical determinant of metabolic and neurological health, carbohydrate quality influences these outcomes in a sex-specific manner.

## Introduction

The prevalence of metabolic disorders such as obesity and type 2 diabetes has dramatically increased over the past several decades ^1,2^. Beyond their direct consequences, obesity and type 2 diabetes significantly increase the risk of other age-related diseases, including Alzheimer’s disease (AD) ^3-9^. Therefore, the need to find effective and affordable ways to improve metabolic health is critical to maintaining the health of an aging population.

Dietary protein is typically thought of as a beneficial nutrient, promoting satiety and muscle growth, and has shown benefits on weight control from increased protein intake in some human trials ^10-14^. In contrast, long-term human studies have shown that protein consumption is associated with an increased risk of diabetes and other age-related diseases ^15-19^. In agreement with these results suggesting excess protein can be detrimental, multiple randomized clinical trials have shown that protein restriction (PR) promotes metabolic health, reducing fat mass, adiposity, and fasting blood glucose, and improving insulin sensitivity, including in type 2 diabetics ^20-23^. These results are recapitulated in animals, with LP diets clearly improving metabolic health, promoting healthy aging and longevity while combatting AD pathology and cognitive impairment ^24-33^. These improvements have been shown to have sex-specific effects in mice, with males exhibiting a more robust response to LP than females ^24,30,34^.

An advantage of studying nutrition in animals is that the dietary composition can be precisely controlled. Prior mouse studies of LP diets have primarily kept diets isocaloric by increasing levels of starch-derived complex carbohydrates; in contrast, modern human diets tend to include readily digestible carbohydrates such as simple sugars, which include the disaccharide sucrose and monosaccharides such as glucose and fructose (commonly in the form of high-fructose corn syrup) ^35^. The consumption of these highly processed carbohydrates has been associated with increased obesity and cardiovascular disease ^36-40^. Fructose is highly lipogenic in itself and also promotes lipogenesis from glucose ^40-43^; high fructose intake has also been implicated in worsened AD pathology in both humans and animal models ^44-49^.

How carbohydrate quality impacts the response to LP diets has been relatively unexplored, but a study by Wali and colleagues assessing how carbohydrate type and digestibility impact the response to dietary protein dilution suggests that this is an important factor in metabolic health^50^. They found that a diet with 10% of calories derived from protein and 70% of calories derived from carbohydrates was the healthiest metabolically when the carbohydrate was comprised of resistant starch but was the worst for health when the carbohydrate source was a 50/50 mixture of fructose and glucose, akin to high-fructose corn syrup. Understanding how dietary carbohydrate quality influences the response to LP diets is critical, as in the real-world, people are free to choose from diverse carbohydrate sources, and replacing protein calories with the “wrong” carbohydrates could potentially lead to negative health outcomes.

Here, we have built on this foundation by comprehensively investigating the metabolic effects of a LP (7% protein) diet when paired with one of four distinct carbohydrate sources: glucose, fructose, a 1:1 glucose/fructose mixture, or sucrose. We examine the metabolic effects of feeding these diets to both male and female C57BL/6J mice. In addition, we examined the impact of protein intake and carbohydrate content on AD pathology and cognition in female APP/PS1 mice. Previous work has shown an effect of amino acid restriction on AD pathology^30,31,51-53^, including in APP/PS1 mice, so we utilized this model to focus on the impact of protein and carbohydrate intake on amyloid pathology. Regarding metabolic health, we find that LP diets are an effective intervention regardless of carbohydrate content, particularly in male C57BL/6J mice. We also found that a fructose-only diet resulted in improved health in almost all metrics measured. In APP/PS1 mice, we found that the metabolic benefits of LP diets are blunted compared to C57BL/6J, but that LP diet reduced amyloid burden. Interestingly, we find that the response to LP diets in female mice is defined by carbohydrate content, with C57BL/6J female mice in the current study exhibiting a greater response to LP diets under certain carbohydrate contexts than in previous studies. These findings have important implications for the translation of LP diet into humans, as the efficacy of LP diets in human studies may also be dependent on other dietary factors.

## Materials and Methods

### Animals

All procedures were performed in accordance with institutional guidelines and were approved by the Institutional Animal Care and Use Committee (IACUC) of the William S. Middleton Memorial Veterans Hospital (Madison, WI, USA) and the University of Wisconsin-Madison IACUC. All mice were maintained at a temperature of approximately 22°C, with a 12:12 light/dark cycle with food and water available *ad libitum*. Health checks were completed on all mice daily. Mice were acclimatized to our experimental facility for one week before experiment start and were housed 2-3 per cage.

For the first experiment, male and female C57BL/6J mice were obtained from The Jackson Laboratory (Stock #000664). At 9 weeks of age, mice were randomly assigned to receive one of eight diets: Sucrose Control ( Suc Con; sucrose as the sole carbohydrate source, 21% of calories from protein), Sucrose LP ( Suc LP; sucrose only, 7% of calories from protein), Glucose/Fructose Control (Gluc/Fruc Con; 50/50 mix of glucose and fructose, 21% protein), Glucose/Fructose LP (Gluc/Fruc LP; 50/50 glucose/fructose, 7% protein), Glucose Control (Gluc Con; glucose only, 21% protein), Glucose LP (Gluc LP; glucose only, 7% protein), Fructose Control (Fruc Con; fructose only, 21% protein), and Fructose LP (Fruc LP; fructose only, 7% protein). All diets were obtained from Inotivo (formerly Envigo). Full diet descriptions, compositions, and item numbers are provided in **Table S1**. Randomization of mice to groups was performed at cage level to ensure all groups had approximately the same starting weight and body composition. Mice were maintained on these diets for 20 weeks.

For the second experiment, male and female APP/PS1 breeding pairs were originally obtained from The Jackson Laboratory/Mutant Mouse Resource and Research Centers (Stock # 034829) and were bred to produce experimental cohorts. At 4 months of age, female APP/PS1 mice were randomly assigned to one of the eight diets described above. Randomization of mice to groups was performed at cage level to ensure all groups had approximately the same starting weight and body composition. Mice were maintained on these diets for 24 weeks.

### In vivo procedures

Food consumption in home cages was measured by moving mice to clean cages, filling the hopper with a measured quantity of fresh diet, and measuring the remainder 3-4 days later. The amount of food consumed was adjusted for the number of mice in each cage, the number of days that had passed, and the relative weights of the mice.

Mouse body composition was analyzed at several points throughout the study using an EchoMRI Body Composition Analyzer (EchoMRI, Houston, TX, USA). Metabolic parameters (O_2_, CO_2_, food composition, respiratory exchange ratio (RER), and energy expenditure) and activity tracking were analyzed using a Columbus Instruments Oxymax/CLAMS metabolic chamber system (Columbus Instruments, Columbus OH, USA). Mice were acclimated to the chamber cages for ∼24 hours and data from a subsequent, continuous 24-hour period was recorded and analyzed.

Glucose tolerance tests were performed by fasting the mice overnight for 16 hours then injecting glucose (1 g kg^-1^) intraperitoneally (I.P.) ^54,55^. Insulin tolerance tests were performed by fasting the mice for 4 hours then injecting insulin I.P. (0.75 U kg^-1^). Blood glucose measurements were taken using Bayer Contour and Bayer Contour Next blood glucose meters (Bayer, Leverkusen, Germany) and test strips. P-407 assays were performed by fasting the mice overnight for 16 hours then re-feeding for 4 hours then injecting P-407 (1 g kg^-1^; donated by BASF, Florham Park, NJ, USA) I.P. and collecting blood at 0, 30, 60, 120, and 180 minutes into microvette tubes containing EDTA (Sarstedt, Inc, Newton, NC, USA). Tubes were then centrifuged at 2000×g for 5 minutes and plasma collected and flash frozen for later analysis. Lipid tolerance tests were performed by fasting mice overnight for 16 hours then administering 2 g kg^-1^ olive oil (Sigma Aldrich, St. Louis, MO, USA) via oral gavage. Blood was collected into microvette tubes containing EDTA at 0, 1, 3, and 5 hours after gavage. Plasma was separated and flash frozen.

Behavioral assays were performed on all APP/PS1 mice and recorded/analyzed using the Ethovision XT Animal Tracking Software by Noldus. Before each procedure mice were allowed to acclimate to the behavioral testing room for 30 minutes. Between each run the arenas and objects were cleaned with 70% ethanol to remove olfactory cues.

For dark/light box, mice were placed into the dark portion of a box that was 1/3 enclosed and 2/3 open, with a small opening for movement between the two zones. The mice were allowed to explore the box for 5 minutes, during which time latency to entry into light zone was measured.

For Novel Object Recognition (NOR), mice were first habituated to the empty arena with no objects. Approximately 24 hours later, an acquisition trial was performed where the mice were placed in the arena with two of the same object, equidistant from the mouse, hereafter referred to as “Object A”. The mouse was allowed to explore the two objects for 5 minutes before returning to the home cage. One hour after the acquisition trial, a short-term memory (STM) test was performed during which the mouse was placed into the arena with one of object A as well as a single novel object, or “Object B”, and allowed to explore for 5 minutes. The following day, 24 hours after the STM test, a long-term memory (LTM) test was performed. During the LTM test, the mouse was placed in the arena with one of object A and a single novel object, “Object C”, and allowed to explore for 5 minutes. Results were quantified by calculating discrimination index (DI) by dividing the time spent exploring the novel object by the time spent examining both objects. A DI value between -0.2 and 0.2 indicates no difference in time spent examining each object. A positive value indicates greater time was spent examining the novel object whereas a negative value indicates greater time was spent examining the familiar object. Due to the nature of the DI calculation, absolute DI is also calculated to separate discrimination from object preference.

For Barnes maze, the test involved 4 phases: habituation, acquisition, and STM and LTM memory tests. During habituation, the mice were placed on the maze and immediately led to the escape box where they remained for 2 minutes before returning to their home cage. After approximately 15 minutes, the mice underwent their first acquisition trial in which they were allowed to explore the maze until either they found the escape box or 3 minutes had elapsed. If the mouse did not find the escape box in 3 minutes, they were gently led to the escape box. Regardless of how the mouse reached the escape box, they remained there for 1 minute after every trial. Over the first four days, four acquisition trials were performed each day with an inter-trial interval of 15 minutes between each trial. On days 5 (STM test) and 12 (LTM test), the escape box was removed and the mouse was placed on the maze for 90s to determine how well it was able to recall where the escape box was. Latency to goal was quantified for each run, and a latency of 180 seconds was assigned to any training runs where the mouse didn’t reach the escape hole while a latency of 90 was assigned for unsuccessful test runs.

Mice were euthanized by cervical dislocation after an overnight fast and a 4-hour re-feed and tissues for molecular analysis were flash-frozen in liquid nitrogen and stored at -80°C or fixed and prepared as described below. For adipose tissue histology, a portion of the inguinal white adipose tissue (iWAT) was collected and fixed in 10% neutral buffered formalin (NBF) for 24 hours before transferring to 70% ethanol for storage until later analysis. For liver histology, a small section of the large lobe was removed and placed in Tissue-Tek Optimal Cutting Temperature (OCT) compound (Sakura Finetek USA Inc, Torrance, CA, USA) and frozen on dry ice before storage at -80°C for later analysis. Brains from APP/PS1 mice were collected. One hemisphere was flash-frozen while the other was fixed in 10% NBF for 48 hours before transferring to 70% ethanol for storage until later analysis.

### Liver and Plasma Assays

Plasma levels of FGF21 were assessed using the Mouse/Rat FGF21 ELISA from R&D Systems (Minneapolis, MN, USA). Plasma and liver concentration of triglycerides and cholesterol were assessed using colorimetric assays from Pointe Scientific (Canton, MI, USA). To assess liver triglyceride and cholesterol levels, livers were homogenized in 95% ethanol. Plasma AST and ALT levels were assessed using ELISAs from Abcam (Cambridge, UK).

### RT-qPCR

To determine expression of metabolism-related genes in iWAT, brown adipose tissue, and liver, RT-qPCR was performed. RNA was extracted using TRI reagent (Sigma Aldrich, St. Louis, MO, USA) according to manufacturer’s protocol and concentration and purity of isolated RNA determined by Nanodrop (Thermo Fisher Scientific, Waltham, MA, USA). 1µg of RNA was used to generate cDNA using Superscript III (Invitrogen, Carlsbad, CA, USA). Oligo dT primers for cDNA synthesis and primers for qPCR were obtained from Integrated DNA Technologies (IDT; Coralville, IA, USA) and are listed in **Table S2**. qPCR reactions were run on a StepOne Plus machine (Applied Biosystems, Foster City, CA, USA) or a QuantStudio qPCR system (Thermo Fisher Scientific, Waltham, MA, USA) using Sybr Green PCR Master Mix (Invitrogen, Carlsbad, CA, USA). β-actin was used to normalize the results from gene-specific reactions.

### Histology

Formalin-fixed iWAT samples and OCT-embedded liver samples were sent to the UW Carbone Cancer Center Experimental Animal Pathology Laboratory (UWCCC EAPL) for further processing. iWAT samples were paraffin embedded then sectioned and H&E stained. Liver samples were cryosectioned and Oil Red O stained.

After sectioning and staining of the samples by UWCCC EAPL, we obtained images of the sections using an EVOS microscope (Thermo Fisher Scientific, Waltham, MA, USA) at 40x magnification as previously described ^56,57^. Three images each from sections per slide were taken, resulting in a total of 9 images per biological replicate. Adipocyte (for iWAT) and lipid droplet (for liver) size was quantified using ImageJ (NIH, Bethesda, MD, USA) and an average value obtained for each biological replicate.

Formalin-fixed half brain samples were paraffin-embedded by the UWCCC EAPL. 5µm sections were then cut using a Leica microtome (Wetzlar, Germany) and affixed to Superfrost Plus microscope slides (Thermo Fisher Scientific, Waltham, MA, USA). For amyloid plaque staining, brain sections were deparaffinized and rehydrated, then treated in 70% formic acid for epitope retrieval. Sections were blocked with normal goat serum (NGS) and incubated with 6E10 monoclonal antibody (1:100; #803004; Biolegend, San Diego, CA, USA) overnight before incubation with biotinylated secondary antibody and visualization with diaminobenzidine chromagen. Glial and amyloid co-staining was performed by deparaffinizing and rehydrating sections followed by antigen retrieval using a 1x Tris-EDTA (pH9.0) antigen retrieval buffer (Abcam, Cambridge, UK) and stained (1:1000) with anti-Iba1 (ab178847; Abcam, Cambridge, UK) and anti-GFAP (PIMA512023; Thermo Fisher Scientific, Waltham, MA, USA) antibodies. Secondary staining was done using a cocktail containing Alexa Fluor 488 goat anti-mouse (1:200; Thermo Fisher Scientific, Waltham, MA, USA), Alexa Fluor 594 goat anti-rabbit (1:200; Thermo Fisher Scientific, Waltham, MA, USA), MethoxyX04 (20µg/mL; Tocris Bioscience, Bristol, England), and To Pro 3 (1:1000, Thermo Fisher Scientific, Waltham, MA, USA). Antibody vendors, catalog numbers and the dilution used are provided in **Table S3**. All sections were imaged using an EVOS FL Auto microscope (Thermo Fisher Scientific, Waltham, MA, USA) at 20x magnification. ImageJ was used for quantification by converting images to binary images via an intensity threshold and quantifying positive area.

### Immunoblotting

Left brain hemisphere samples were lysed in cold RIPA buffer supplemented with phosphatase inhibitor and protease inhibitor cocktail tablets (Thermo Fisher Scientific, Waltham, MA, USA) using a FastPrep 24 (M.P. Biomedicals, Santa Ana, CA, USA) with bead-beating tubes from (VWR, Radnor, PA, USA) and zirconium ceramic oxide beads from (Thermo Fisher Scientific, Waltham, MA, USA). Protein lysates were then centrifuged at 13,300 rpm for 10 min and the supernatant was collected. Protein concentration was determined by Bradford Assay (Pierce Biotechnology, Waltham, MA, USA). 20-60 μg protein was separated on 8%, 10%, or 16% tris-glycine gels (ThermoFisher Scientific, Waltham, MA, USA) and transferred to PVDF membrane (EMD Millipore, Burlington, MA, USA). Autophagy markers were assessed, including autophagy proteins Atg5, Atg7, and Atg16L1, autophagosome formation proteins Beclin and LC3A/B, and the autophagy receptor (sequestosome 1, SQSTM1). Antibody vendors, catalog numbers and the dilution used are provided in **Table S3**. Imaging was performed using a Bio-Rad Chemidoc MP imaging station (Bio-Rad, Hercules, CA, USA). Quantification was performed by densitometry using NIH ImageJ software.

### Statistical Analyses

Z-scores were calculated using the mean and standard deviation of the pooled control population for male and female mice separately. All statistical analyses were conducted using Prism, version 10 (GraphPad Software Inc., San Diego, CA, USA). Tests involving multiple factors were analyzed by a three-way analysis of variance (ANOVA) with carbohydrate type, protein level, and sex as variables. To perform multiple comparisons, a two-way ANOVA was performed for each metric with carbohydrate type and protein level as variables, followed by a Dunnett’s, Tukey-Kramer, or Sidak’s post-hoc test as specified in the figure legends. Results of two-way ANOVAs and exact p-values of multiple comparisons are provided in **Table S4**.

## Results

### Both low protein and fructose-only diets improve body composition

Male and female 10-week-old C57BL/6J mice were fed diets containing either 21% (Control) or 7% (Low Protein; LP) protein, and one of 4 different carbohydrate compositions: sucrose, a 1:1 glucose/fructose mixture, glucose, or fructose, for a total of 8 different diets (**Fig. 1A**). All diets were isocaloric, with carbohydrates increasing to replace protein calories in the LP diets; all diets had identical fat composition. The complete composition of all diets is detailed in **Table S1**.

**Figure 1:**
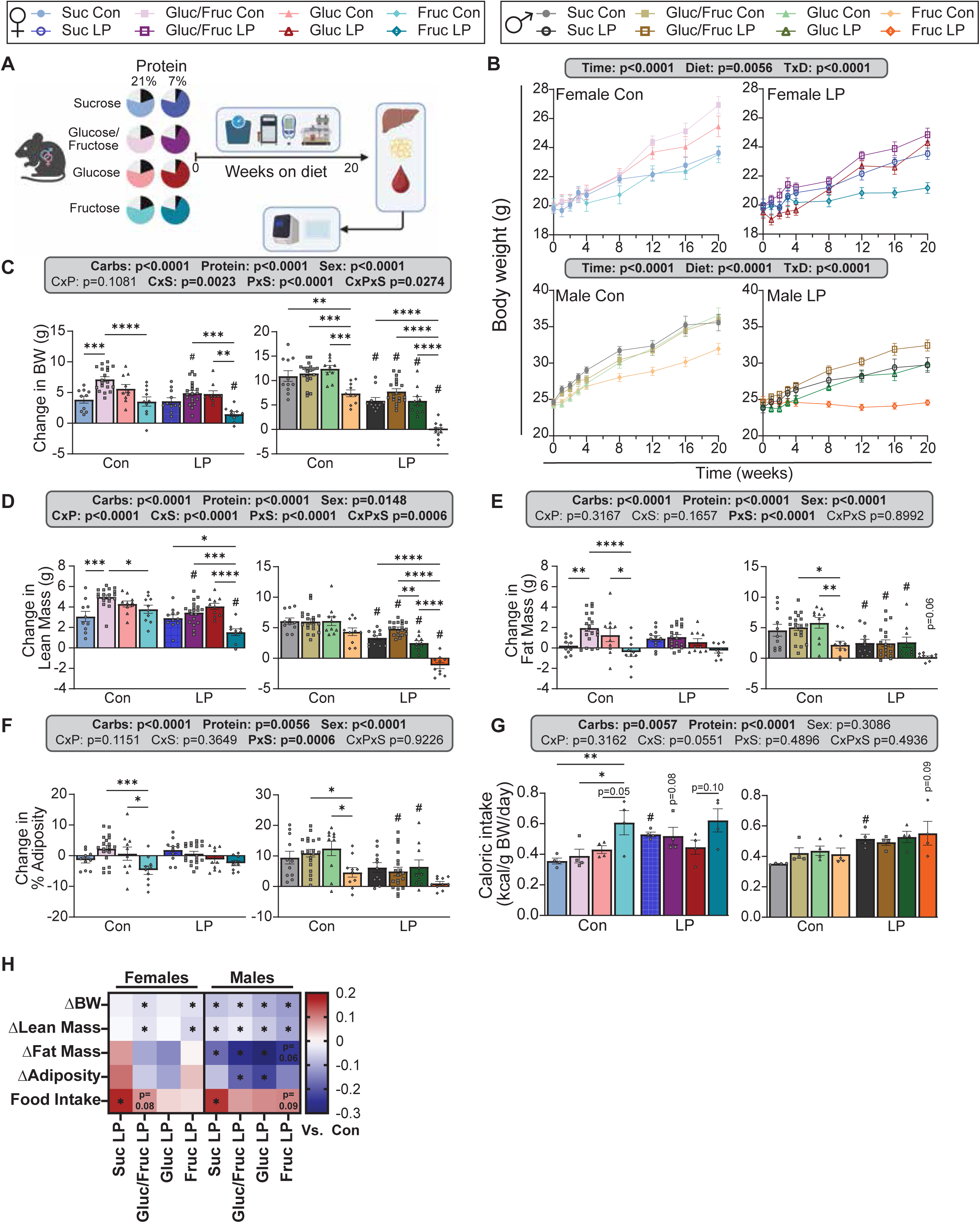
Both LP and fructose-only diets promote leanness in both female and male mice. (A) Experimental schematic; female and male C57BL/6J were maintained on control and LP diets containing sucrose, glucose/fructose, glucose, or fructose for 20 weeks starting at 9 weeks of age before euthanasia and tissue collection. (B) Female and male body weights over the 20-week experiment (B) and overall change in body weight at the end of the experiment (C). (D-F) Body composition was measured using an EchoMRI Body Composition Analyzer to determine final lean mass (D), fat mass (E), and adiposity (F). (G) Home cage food consumption was determined and normalized to body weight. (H) Heat map representing log_2_ fold change body weight, composition, and food consumption changes in response to LP diets. Asterisks indicate metrics that exhibited significant differences between control and LP within each carbohydrate type. (A-G) Body weight and composition data n=10-19/group; home cage data n=4/group. Asterisks denote differences between carbohydrate types within control or LP diets with *p<0.05, **p<0.01, ***p<0.001, ****p<0.0001, Tukey test post 2-way ANOVA. Hashes denote significant differences (p<0.05, Tukey test post 2-way ANOVA) in LP diets compared to their corresponding control counterparts. Exact p-values are provided in Table S3.

Over the course of 20 weeks, LP-fed mice of both sexes gained less weight than their Con-fed counterparts (**Figs. 1B-C**). As we expected ^24,58^, male mice responded more strongly to LP than females, but the magnitude of this difference was dependent on carbohydrate type. Male mice fed diets with fructose as the carbohydrate source had blunted body weight accretion compared to all other carbohydrate sources, irrespective of their consumption of a Control or LP diet; a similar effect of fructose on weight accretion was seen in females, especially in contrast with Gluc/Fruc-fed females.

We determined the effects of each diet on body composition. There was an overall effect of both carbohydrate source and protein intake on lean mass in both sexes (**Figs. 1D, S1A-B**). In females, LP reduced lean mass only in diets containing fructose monosaccharides, and females fed a fructose-only diet had the lowest lean mass in both control and LP contexts. Interestingly, while Fruc LP-fed female mice simply had blunted lean mass accretion, Fruc LP-fed male mice actually lost lean mass over the course of the experiment. LP diet reduced lean mass accretion compared to Con in male mice in all carbohydrate contexts.

Changes in fat mass (**Figs. 1E, S1C-D**) and adiposity (**Figs. 1F, S1E-F**) exhibited more sexual dimorphism, with females responding very strongly to carbohydrate type but not protein content, while males responded to both. In females, both sucrose and fructose diets reduced fat mass as compared to other Con diets, but only mice fed the Fruc Con diet had significantly decreased adiposity. There was no significant effect of carbohydrate type on the fat or adiposity of LP-fed females. In contrast, Con-fed males had the lowest fat mass and adiposity when consuming a fructose only diet. LP diet decreased fat mass and adiposity in all males regardless of carbohydrate type, with the effects on adiposity reaching significance only in males consuming LP diets with glucose monosaccharides.

Finally, there was an overall positive main effect of an LP diet on food intake in both sexes that reached statistical significance when sucrose was the carbohydrate source (**Fig. 1G**). In females, fructose consumption increased food intake regardless of dietary protein content, reaching statistical significance against several other carbohydrate sources. Despite the stimulatory effect of LP diet on food intake, LP-fed mice still exhibited overall reduced protein intake and increased carbohydrate intake (**Figs. S1G-H**).

In conclusion, female mice had a more limited response to LP diets, with the response varying by carbohydrate type, while the response to LP diets in males was stronger. The ability of carbohydrate type to impact the response to LP diet was less striking in males than in females (**Fig. 1H**).

### Low protein diets promote beiging and thermogenic gene expression in white adipose tissue in male mice

The reduced weight and adiposity of LP-fed mice despite their increased caloric intake, is partially driven by increased energy expenditure (EE) (**Figs. 2A-B**). In females, LP diets significantly increased EE during both the light and dark period when the diets contained glucose monosaccharides, and during the dark period when the diet contained sucrose. In contrast, LP-fed females consuming a fructose diet did not have a statistically significant increase in EE relative to Con-fed females, and Fruc LP-fed females had reduced EE relative to Gluc LP-fed (light cycle) and Suc LP-fed (dark cycle) females. Males fed LP diets had increased EE in both the light and dark periods in all but the sucrose diets. Gluc Con-fed male mice had reduced EE compared to other Con diets in both the light and dark periods while Fruc LP-fed male mice had increased EE compared to Suc LP-fed mice during the light period.

**Figure 2:**
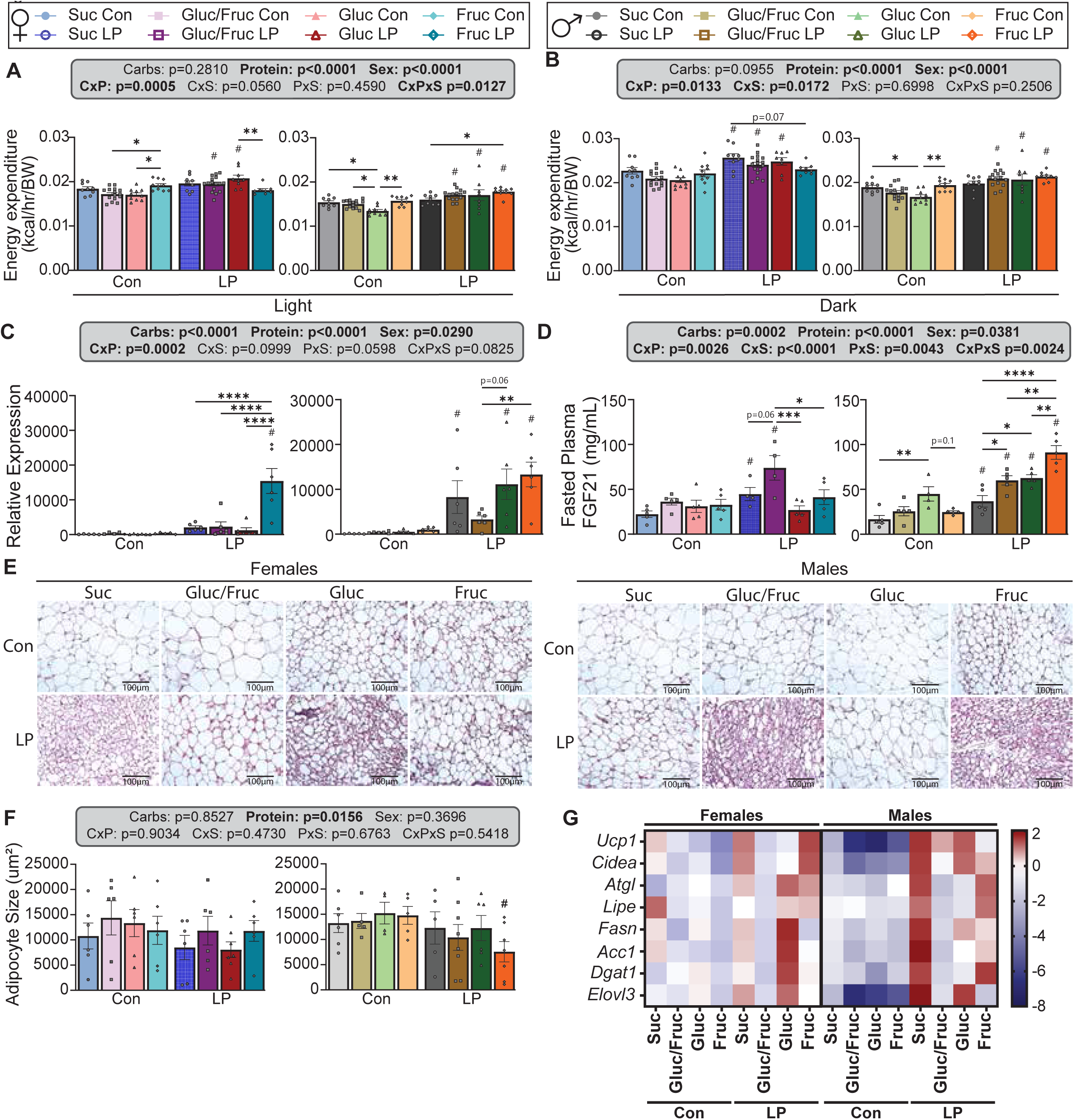
LP diet increases energy expenditure, FGF21 levels, iWAT beiging, and iWAT thermogenic gene expression. (A-B) Female and male mice were placed into Columbus Instruments metabolic chambers/CLAMS to determine energy expenditure during the light (A) and dark (B) periods. n=10-19 (C-D) *Fgf21* levels in female and male mice were determined in liver (C) as well as plasma levels of FGF21 (D). n=4-5 (E-F) H&E staining in iWAT was used to determine beiging and adipocyte size in female and male mice. Representative images are shown in (E) and adipocyte size quantified in (F). (G) qPCR was performed to determine expression of thermogenic, lipolytic, and lipogenic-related genes in iWAT in both female and male mice. Results are shown in a heat map compared to the average of all groups within each sex. n=4-5. All comparisons in this figure: asterisks denote differences between carbohydrate types within control or LP diets with *p<0.05, **p<0.01, ***p<0.001, ****p<0.0001, Tukey test post 2-way ANOVA. Hashes denote significant differences (p<0.05, Tukey test post 2-way ANOVA) in LP diets compared to their corresponding control counterparts. Exact p-values are provided in Table S3.

Respiratory exchange ratio (RER) is calculated from the ratio of O_2_ consumed and CO_2_ produced; a value close to 1.0 indicates that carbohydrates are primarily utilized for energy production, while a value approaching 0.7 indicates that lipids are the predominant energy source ^59,60^. As we substituted carbohydrates for protein in the LP diets, we expected RER to increase in LP groups. Indeed, LP broadly increased RER during the dark cycle in both sexes (**Figs. S2A-B**). During the light cycle, RER was increased significantly only in Gluc LP-fed mice. Carbohydrate quality further modulated RER, especially in Con-fed male mice, with Gluc Con-fed males having lower RER and Fruc Con-fed males having the highest, reaching significance when compared against each other as well as when compared to Gluc/Fruc Con-fed males during both the light and dark periods.

Low protein diets increase energy expenditure by inducing hepatic production of FGF21 ^25,28,61^. As expected, LP feeding broadly upregulated hepatic *Fgf21,* with a stronger effect in males than females (PxS, p=0.0598), but we also observed an overall significant effect of carbohydrate as well as a significant carb x protein interaction (**Fig. 2C**). In females, the induction of *Fgf21* by LP was restricted to the Fruc LP-fed mice, while in males the effect was significant in Suc LP, Gluc LP, and Fruc LP-fed mice. Plasma FGF21 was significantly induced by LP in females in the Suc and Gluc/Fruc-fed mice, while in males all LP diets elevated FGF21, with Fruc LP-fed males having significantly higher plasma levels than all other groups (**Fig. 2D**). The hepatic expression of *Fgf21* was only partially reflected in plasma levels of FGF21, likely due to differences between the feeding state of the liver mRNA (refed) and plasma (fasted).

To determine if FGF21 induction induced white adipose tissue (WAT) thermogenesis, we examined the inguinal WAT (iWAT) depot. We observed an overall effect of protein on adipocyte size, with LP-fed males and females having smaller adipocytes, though the effect for any given diet was significant only in Fruc LP-fed males (**Figs. 2E-F**). RT-qPCR analysis of thermogenic gene expression showed that LP increased *Ucp1* and *Cidea* expression in both sexes, reaching significance in Fruc LP-fed females and Suc LP and Gluc LP-fed males (**Figs. 2G, S2E-F**). LP also induced genes related to lipolysis and lipogenesis, suggesting an increase in futile lipid cycling (**Figs. 2G, S2F-L**). In females, carbohydrate-specific effects were limited to the LP context, where the Gluc LP diet uniquely upregulated *Fasn* and *Dgat1* (**Figs. S2I, K**). In males, the LP diet broadly upregulated *Atgl*, *Lipe*, *Fasn*, *Acc1*, and *Dgat1*, with Suc LP and Fruc LP-fed males showing increased expression of *Atgl*, *Lipe*, and *Dgat1* (**Figs. S2G, H, K**), and Suc LP-fed males had increased *Fasn* and *Acc1* (**Figs. S2I-J**).

Overall, we found that as expected, LP diets stimulate EE, inducing the FGF21-UCP1 axis and iWAT beiging. Importantly though, these responses are highly dependent upon both sex and, especially in females, carbohydrate source.

### Glucose homeostasis is improved in both female and male mice fed LP diets, whereas lipid homeostasis is more dependent on carbohydrate type

LP diets broadly decreased fasting blood glucose in both sexes compared to control diets, significantly in the case of Gluc/Fruc LP-fed mice of both sexes and Gluc LP-fed males (**Fig. S3A**). LP diets improved glucose tolerance in all groups of both sexes, irrespective of carbohydrate type (**Figs. 3A-B**). However, carbohydrate type did impact glucose tolerance; Gluc Con and Fruc Con fed females had better glucose tolerance relative to other Con-fed females; meanwhile, Fruc-fed males had the best glucose tolerance in both Con and LP contexts and Gluc LP-fed males had superior glucose tolerance than Suc LP-fed males (**Figs. 3A-B**). In contrast, sensitivity to IP insulin was largely unaffected by LP feeding, with LP enhancing insulin sensitivity only in Gluc-LP fed males (**Figs. 3C-D**). Analysis of fasting insulin to perform a homeostasis model assessment of insulin resistance (HOMA-IR) revealed an effect of protein, but not carbohydrates, in females, with improved insulin sensitivity (reduced HOMA-IR) in Fruc LP-fed mice compared to Con (**Fig. S3B**). Males, however, do show an effect of carbohydrates as well, with Gluc/Fruc Con mice having increased insulin resistance, while both Gluc and Gluc/Fruc LP groups had improved insulin sensitivity compared to their control counterparts.

**Figure 3:**
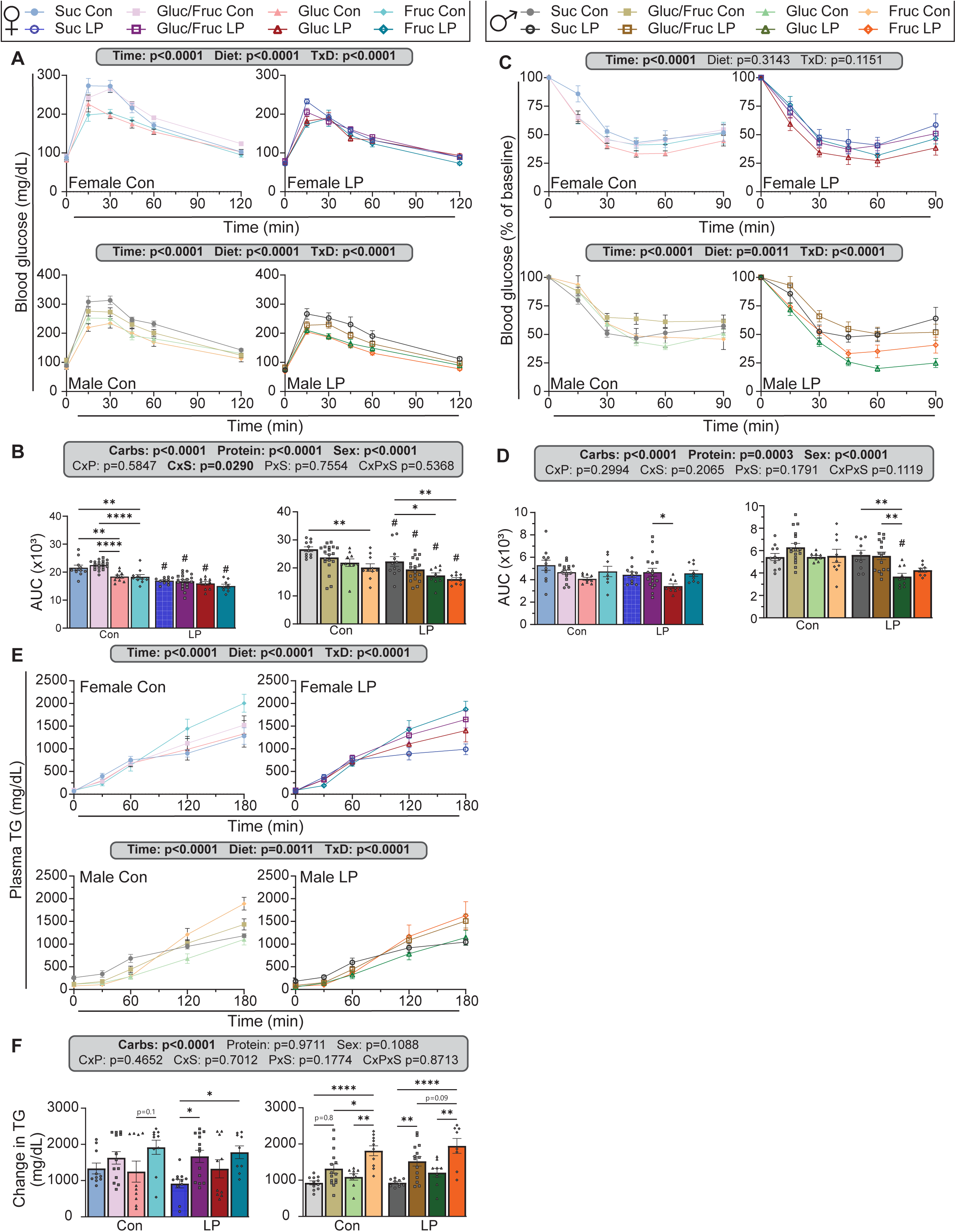
Glucose homeostasis is more strongly impacted by protein intake whereas lipid homeostasis is only affected by carbohydrate type. (A-B) Glucose tolerance tests were performed in female and male (A) and area under the curve quantified (B) to determine glucose tolerance. (C-D) Insulin tolerance tests were performed in female and male (C) and area under curve quantified (D) to determine insulin sensitivity. (E-F) P-407 assays were performed in female and male mice (E) as an *in vivo* approximation of lipid homeostasis and change in TG over the 3 hour assay quantified (F). n=10-19. All comparisons in this figure: asterisks denote differences between carbohydrate types within control or LP diets with *p<0.05, **p<0.01, ***p<0.001, ****p<0.0001, Tukey test post 2-way ANOVA. Hashes denote significant differences (p<0.05, Tukey test post 2-way ANOVA) in LP diets compared to their corresponding control counterparts. Exact p-values are provided in Table S3.

To evaluate triglyceride production, we treated mice with P-407, a lipoprotein lipase inhibitor that blocks peripheral lipid uptake and thus enables the quantification of triglyceride production ^62^. Interestingly, there was no overall effect of protein on plasma triglyceride (TG) levels during this assay (**Figs. 3E-F**). Instead, the primary driver of differences between groups was carbohydrate type; both sexes had significantly higher plasma TGs when consuming diets containing fructose monosaccharides. Plasma cholesterol levels showed a similar driving role for carbohydrates during the assay (**Figs. S3C-D**).

Finally, we performed a lipid tolerance test. We found a significant overall interaction between carbohydrate type, protein, and sex (**Figs. S3E-F**). In females, Fruc Con-fed mice had the fastest lipid clearance, whereas diets containing glucose monosaccharides had the most rapid clearance in the LP context. An LP diet improved clearance relative to the Con diet of the same carbohydrate type only when the mice consumed the Gluc/Fruc mixture. In sharp contrast to these results in females, we observed no effect of either protein or carbohydrates on lipid clearance in male mice.

These results show a stronger impact of protein intake on glucose homeostasis, while carbohydrate type, particularly the presence of fructose, is the primary driver of lipid homeostasis.

### Dietary carbohydrate type and protein level both regulate liver carbohydrate and lipid metabolism

To investigate the effects of these diets on liver lipids, we performed Oil Red O staining and measured liver triglyceride levels (**Figs. 4A-C**). In females, lipid droplet size was not significantly altered by carbohydrates or proteins, though there was a clear carbohydrate by protein interaction, with droplet size increasing with LP feeding when sucrose was the carbohydrate source and smaller when fructose was the sole carbohydrate (**Figs. 4A-B**). In males, LP diets had an overall effect of increased lipid droplet size, which reached significance only in mice fed diets with fructose monosaccharides. Gluc Con-fed males had larger lipid droplets than other Con-fed groups, but the size of the drops was not increased by LP feeding (**Figs. 4A-B**).

**Figure 4:**
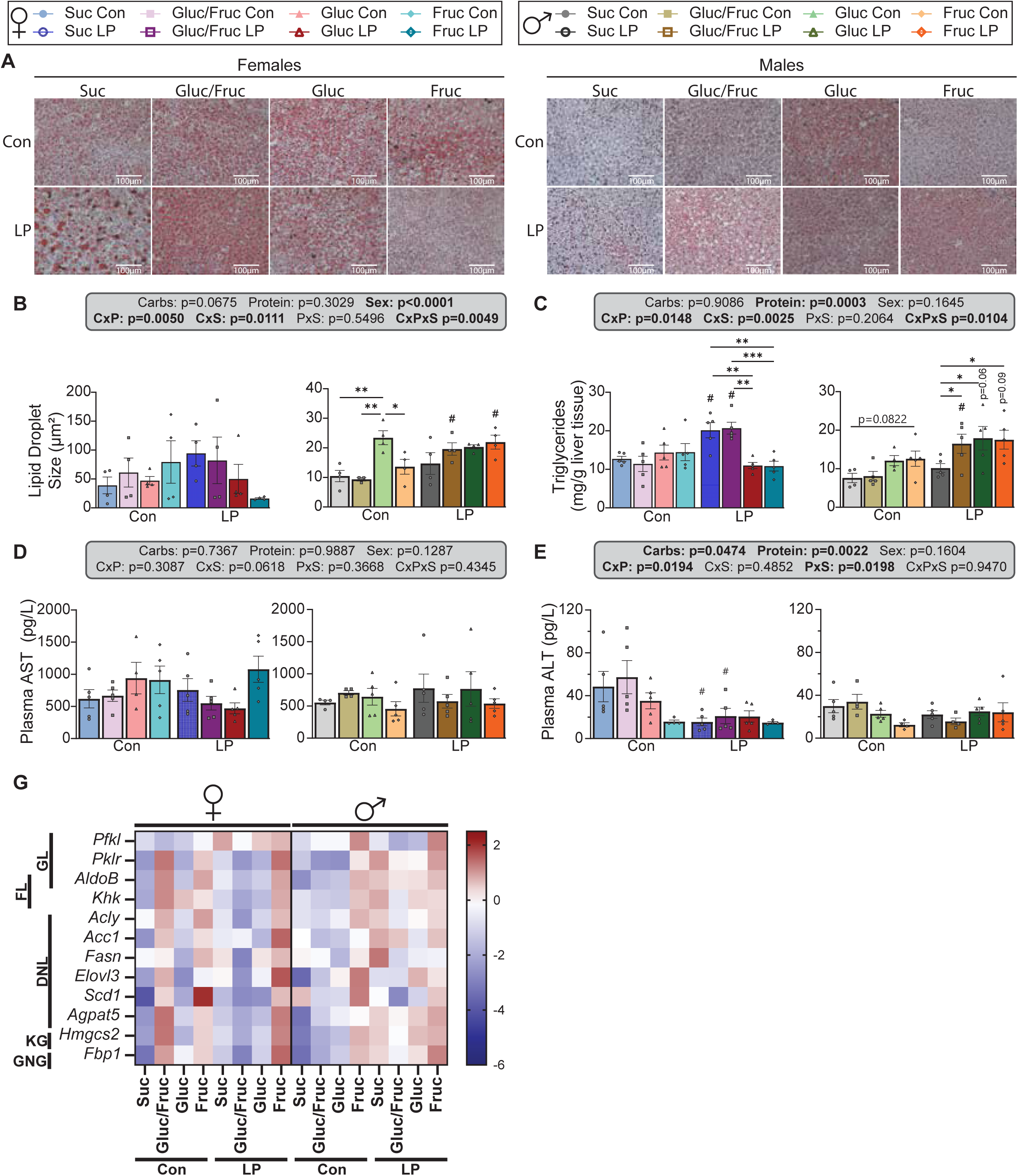
Liver metabolism is impacted by both carbohydrate type and protein intake. (A-C) Lipid droplet deposition and triglyceride levels in female and male mice, as determined by Oil Red O staining (A) and quantification (B) and colorimetric assay for triglycerides (C). (D-E) Plasma AST (D) and ALT (E) levels were quantified to assess liver damage. (F) Gene expression for metabolism-related genes in the liver of female and male mice. Results are shown in a heat map compared to the average of all groups within each sex. n=4-5. All comparisons in this figure: asterisks denote differences between carbohydrate types within control or LP diets with *p<0.05, **p<0.01, ***p<0.001, ****p<0.0001, Tukey test post 2-way ANOVA. Hashes denote significant differences (p<0.05, Tukey test post 2-way ANOVA) in LP diets compared to their corresponding control counterparts. Exact p-values are provided in Table S3.

Hepatic triglyceride levels similarly had a complex interplay, with an overall increase with LP feeding, but strong interactions with carbohydrate type and sex. An LP diet increased triglyceride levels in females only when the carbohydrate source contained both glucose and fructose (**Fig. 4C**). In males, an LP diet failed to increase triglyceride levels only when the mice consumed sucrose (**Fig. 4C**). Although hepatic lipids are associated with hepatic steatosis, circulating markers of liver damage (AST and ALT) suggest an absence of toxicity. Plasma AST levels were not affected at all by diet (**Fig. 4D**). ALT levels showed an overall significant effect of protein and a protein by sex interaction, which we interpret as ALT decreasing in LP-fed females, which was statistically significant in the case of Suc LP and Gluc/Fruc LP-fed females (**Figs. 4E**). This suggest that the “fatty liver” induced by LP is non-pathological and may represent a healthy form of lipid storage for rapid utilization.

We examined the molecular drivers of these changes by examining the expression of fructose, glucose, and lipid metabolism genes in the liver (**Figs. 4G, S4**). While almost every gene showed an effect of sex or an interaction of sex with protein or carbohydrate, we observed that liver metabolism was more strongly influenced by carbohydrate type than protein level. In females, diets containing fructose monosaccharides (Gluc/Fruc and Fruc) generally had increased expression of genes involved in fructose metabolism (*AldoB* and *Khk*; **Figs. 4G, S4C-D**), *de novo* lipogenesis (*Acly*, *Acc1*, *Fasn*, *Elovl3*, and *Scd1*; **Figs. 4G, S4E-I**), and ketogenesis (*Hmgcs2*; **Fig. 4G, S4K**). This effect was most pronounced in Con-fed females, but Fruc LP-fed mice also exhibit increases in most of these same genes. In males, the Fruc Con diet upregulated expression of *Pfkl*, *Acly*, *Elovl3*, *Scd1*, and *Fbp1* (**Figs. 4G, S4A, E, H, I, L**); others genes (*Pklr*, *AldoB*, *Khk*, *Hmgcs2*) showed an overall effect of carbohydrates, which we interpret as likely drive by increased expression in Fruc Con-fed mice.

### Carbohydrate type defines the response to protein restriction in female, but not male, mice

To evaluate the overall effects of dietary carbohydrate and protein on metabolic health, we performed multivariate analyses. We correlated the protein calories consumed by each individual mouse with all the collected phenotypic and molecular metrics, and visualized these with a heat map using hierarchical clustering on the strength and direction of each correlation (**Fig. 5A**). Metrics negatively correlated with protein intake – meaning they are robustly increased in LP-fed mice across sexes and diets – were clustered at the top, and include thermogenic and metabolic gene expression in iWAT, as well as hepatic *Fgf21* expression. Conversely, metrics positively correlated with protein intake (and thus decreased with LP feeding) are clustered at the bottom and include body weight and glucose tolerance. Outlined in black boxes are measures that are consistent within males regardless of carbohydrate type but in females have opposite responses to protein intake depending on dietary carbohydrates. With this, we found that carbohydrate type determines the response of adiposity and liver metabolism in females.

**Figure 5:**
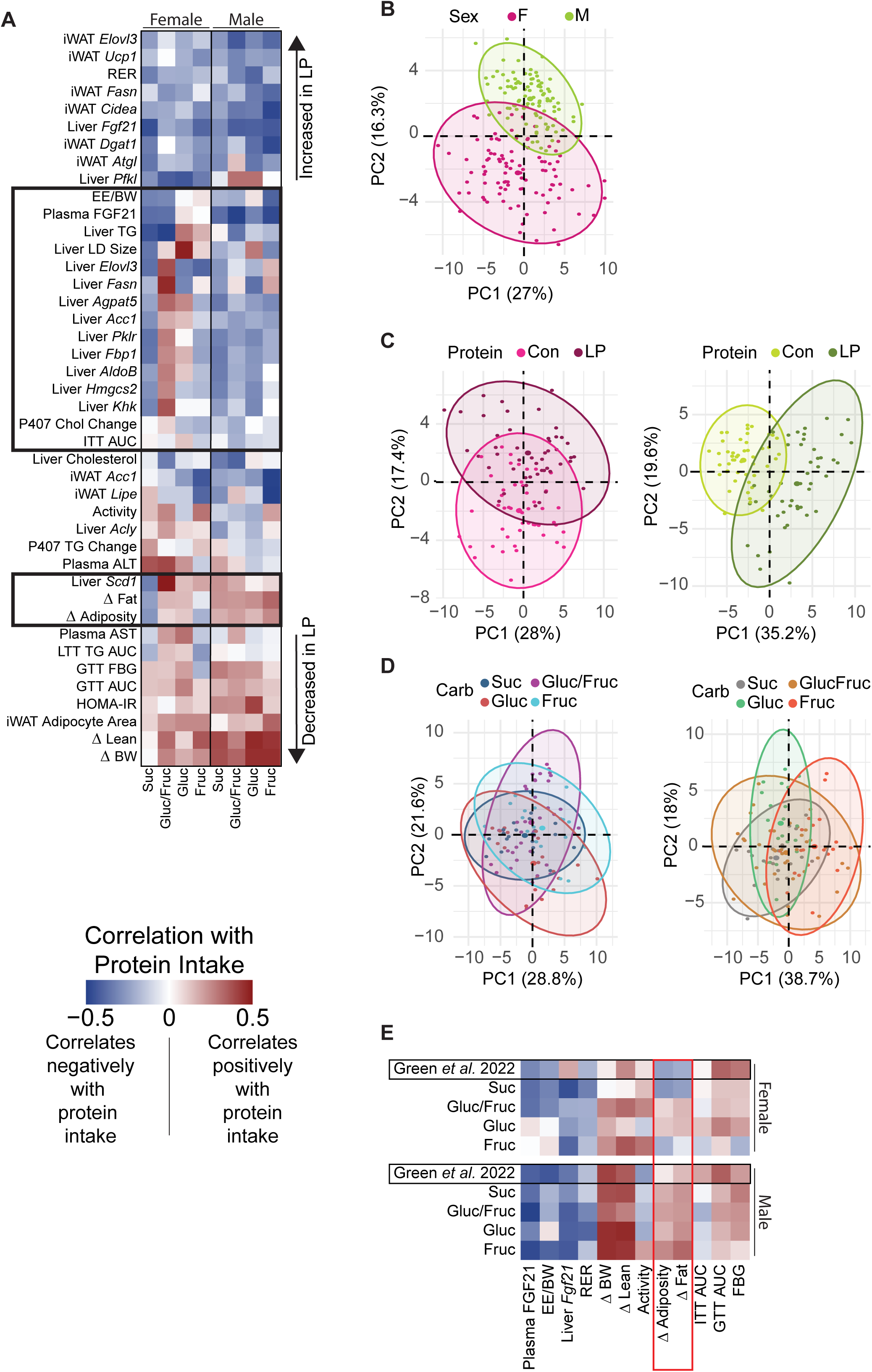
Carbohydrate type modulates the female response to protein restriction. (A) Phenotypic and molecular measures were correlated with protein consumption in each mouse by Pearson’s correlation and clustered by hierarchical clustering. Dark blue indicates measures that correlate negatively with protein intake while dark red indicates measures that correlate positively with protein intake. Black boxes bracket measures that are consistent between carbohydrate contexts in males but not females. (B-D) Individual mice visualized by sex (B), protein intake (C), and carbohydrate intake (D) indicating separation between the groups along Dimension 1 and Dimension 2. (E) Phenotypic measures from the current study and Green *et al.* 2022 ^24^ were correlated with protein consumption in each mouse by Pearson’s correlation and clustered by hierarchical clustering. Dark blue indicates measures that correlate negatively with protein intake while dark red indicates measures that correlate positively with protein intake. The red box brackets measures that are consistent in males, but different in some carbohydrate types in female mice. All comparisons in this figure: asterisks denote differences between carbohydrate types within control or LP diets with *p<0.05, **p<0.01, ***p<0.001, ****p<0.0001, Tukey test post 2-way ANOVA. Hashes denote significant differences (p<0.05, Tukey test post 2-way ANOVA) in LP diets compared to their corresponding control counterparts. Exact p-values are provided in Table S3.

To determine which metrics contribute the most to differences between groups, we performed a principal component analysis (PCA; **Figs. 5B-D, S5A**). We found that the primary contributors to dimensions 1 and 2 across all PCAs were body composition and liver and iWAT gene expression (**Fig. S5A**). Overall, there was a strong effect of sex and protein (**Figs. 5B-C**). In contrast, carbohydrate type alone did not strongly contribute, resulting in overlap between all groups (**Fig. 5D**). This suggests that carbohydrate type does not have consistent effects on all aspects of metabolic health; carbohydrate type has a different magnitude of effect on each aspect of metabolic health examined.

To better examine how sugars alter the response to PR, we re-analyzed our protein correlation data along with previous results using maltodextrin and corn-derived carbohydrates ^24^ from our lab (**Fig. 5E**). We find that male mice in both this historical study and the current one responded very similarly to PR. However, in females, adiposity and fat mass were heavily dictated by carbohydrate type. In the previous study, female adiposity and fat mass were negatively correlated with protein intake, as we also observed in females consuming Suc or Fruc diets. Surprisingly, female mice consuming Gluc/Fruc or Gluc as the dietary sugar respond more similarly to male mice and show a positive correlation of adiposity and fat mass with protein intake.

Together, these results demonstrate that the metabolic health of male mice is primarily driven by dietary protein and respond favorably and consistently to LP diets independent of dietary carbohydrate source. Conversely, in female mice dietary carbohydrate quality and protein interact to determine the metabolic response to LP diets.

### APP/PS1 female mice have worsened metabolic health on diets containing 50/50 glucose/fructose, while LP diets ameliorate this effect

Restriction of protein or the branched-chain amino acids slows the progression and development of Alzheimer’s disease (AD) in mouse models including APP/PS1 mice ^30,31,51-53^. We focused on females as we previously found that LP diets can successfully reduce amyloid load and improve spatial memory in female 3xTg AD mice, and we observed above that in females there is a strong interaction between carbohydrate quality and protein level.

Four-month-old female APP/PS1 mice were placed on the eight previously described diets (**Table S1**) for six months, during which time metabolic and cognitive function were assessed (**Fig. 6A**). Diet-induced changes in body weight, lean and fat mass, and adiposity generally mirrored that seen in wild-type C57BL/6J females, but the magnitude of the response was blunted, with many changes failing to reach statistical significance (**Figs. 6B-F, S6A-C**). However, there was still a strong effect of carbohydrate quality on many phenotypes: APP/PS1 females fed the Gluc/Fruc Con gain more body weight, fat mass, and increased in adiposity more than other control-fed groups (**Figs. 6C-F**).

**Figure 6:**
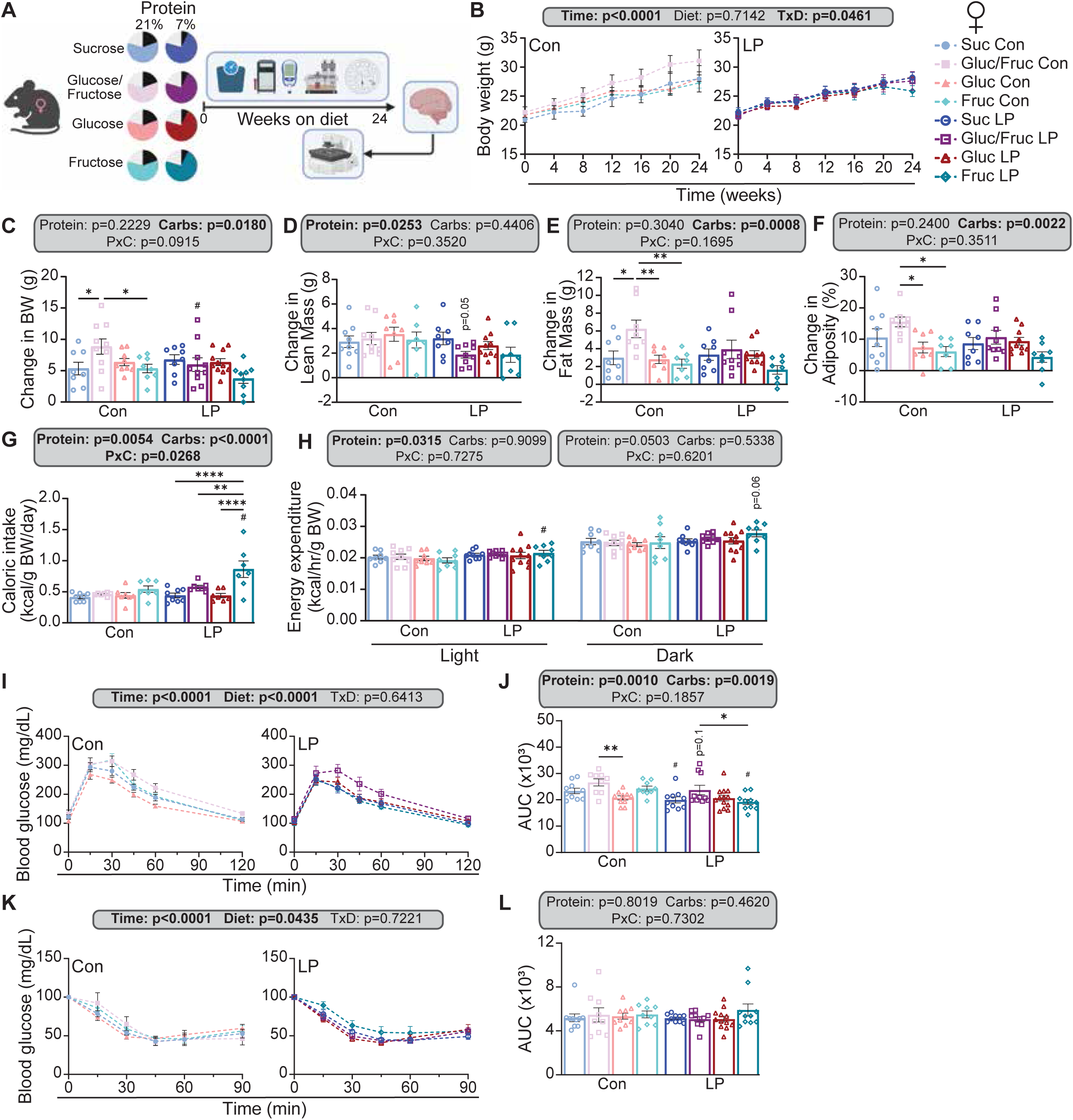
Female APP/PS1 mice exhibit differences in metabolic health in response to both protein and carbohydrate intake. (A) Experimental schematic; female APP/PS1 mice were placed onto one of eight diets containing control or low protein and sucrose, glucose/fructose, glucose, or fructose as carbohydrate source at 4 months until 10 months of age. (B-C) Body weights were measured monthly (B) and final change in body weight (C) calculated. (D-F) Using an EchoMRI body composition analyzer, lean mass (D) and fat mass (E) were measured and adiposity calculated (F). (G) Caloric intake was calculated in home cages and normalized to body weight. (H) Energy expenditure was calculated and normalized to body weight. (I-L) Glucose homeostasis was assessed via glucose (I-J) and insulin (K-L) tolerance tests. Area under the curve was calculated for each test (I,L). n=8-11. All comparisons in this figure: asterisks denote differences between carbohydrate types within control or LP diets with *p<0.05, **p<0.01, ***p<0.001, ****p<0.0001, Tukey test post 2-way ANOVA. Hashes denote significant differences (p<0.05, Tukey test post 2-way ANOVA) in LP diets compared to their corresponding control counterparts. Exact p-values are provided in Table S3.

As in C57BL/6J females, we observed that fructose increased caloric intake, which was significantly higher in Fruc LP-fed mice than other LP-fed groups (**Fig. 6G**). Due to this increase, Fruc Con-fed and Fruc LP-fed mice, consumed both more protein and more carbohydrates than other Con-fed or LP-fed mice, respectively (**Figs. S6D-E**). As expected, RER was increased in LP-fed mice, likely due to the increased carbohydrate content of these diets (**Fig. S6F**). Surprisingly, unlike in C57BL/6J females, we did not find changes in energy expenditure that would account for the increased body weight and adiposity of Gluc/Fruc Con-fed mice (**Fig. 6H**).

AD patients and mouse models exhibit impaired glucose homeostasis ^63^, and we therefore assessed glycemic control. There was a strong effect of protein level on fasting blood glucose, with LP diets decreasing fasting blood glucose, which was significant when fructose was the sole carbohydrate (**Fig. S6G**). Glucose tolerance was robustly impacted by both protein intake and carbohydrate type; LP-fed mice had overall better glucose tolerance, while Gluc/Fruc-fed mice had the worst glucose tolerance in both the Con and LP contexts (**Figs. 6I-J**). All LP diets aside from glucose improved glucose tolerance compared to their control counterparts. The response to I.P. insulin administration was similar in all groups (**Figs. 6K-L**). These effects of protein and carbohydrate are similar, but blunted, relative to those we observed in C57BL/6J females.

### Protein restriction reduces β-amyloid plaque load, while carbohydrate quality impacts spatial memory

We performed cognitive testing at 9 months of age. We found an overall effect of protein (p=0.0166) on movement in the open field test, which we interpret to mean that LP-fed mice had slightly increased motility (**Fig. S7A**). There was no effect of diet on anxiety or object recognition, as determined by dark/light box (**Fig. S7B**) and novel object recognition tests (**Figs. S7C-F**). However, assessment of spatial memory using the Barnes maze identified an effect of carbohydrate quality on short-term memory (**Figs. 7A-C**). Gluc/Fruc-fed mice had the worst STM irrespective of protein level, with Gluc/Fruc Con-fed mice having significantly worse STM performance than Gluc Con or Fruc Con-fed mice (**Fig. 7B**). This is consistent with worsening of metabolic health in Gluc/Fruc-fed mice. There was no effect of diet on performance during training or on long-term memory (**Figs. 7A, 7C**).

**Figure 7:**
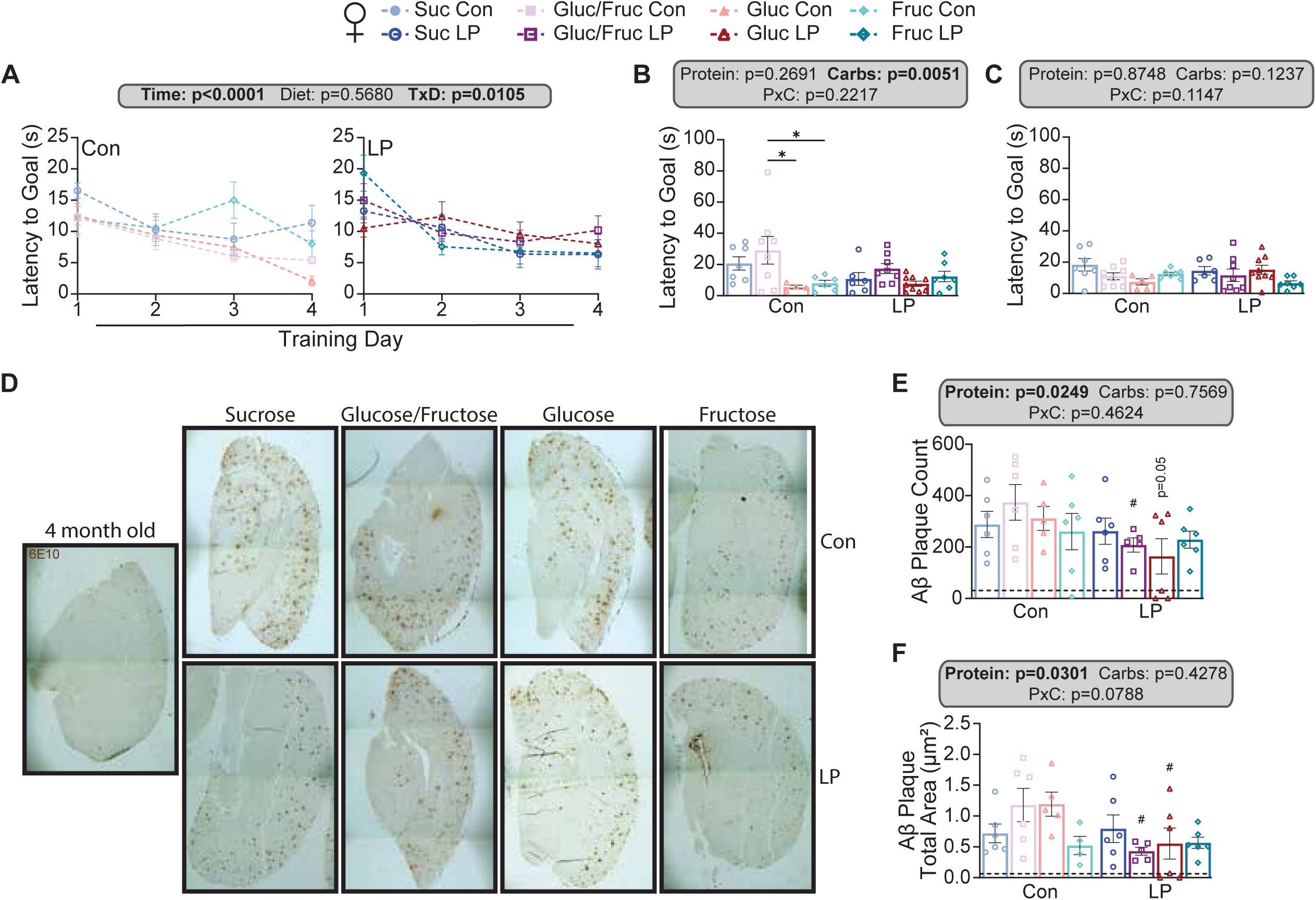
LP diet reduced amyloid burden in APP/PS1 mice but spatial memory was worsened in Gluc/Fruc females only. (A-C) Barnes maze was performed with training over 4 days (A) followed 24 hours later by short-term memory test (B) and a long-term memory test 7 days later (C). (D-F) Amyloid 6E10 DAB staining was performed on the midbrains of APP/PS1 mice. Images were taken on Evos at 4X magnification then stitched to acquire whole midbrain images; dark horizontal lines are an artifact of stitching (D). Quantification of plaque count (E) and total plaque area (F) was performed. Dotted lines indicate levels found in 4-month-old APP/PS1 mice. n=5-9. All comparisons in this figure: asterisks denote differences between carbohydrate types within control or LP diets with *p<0.05, **p<0.01, ***p<0.001, ****p<0.0001, Tukey test post 2-way ANOVA. Hashes denote significant differences (p<0.05, Tukey test post 2-way ANOVA) in LP diets compared to their corresponding control counterparts. Exact p-values are provided in Table S3.

We assessed AD pathology via postmortem staining for β-amyloid (Aβ), microglia and astrocytes (**Figs. 7D-F and S7G-K**). As we expected, there was a significant effect of protein on both plaque count and total plaque area (protein effect p=0.0249 and p=0.0301, respectively), with reduced AD pathology in LP-fed mice. Aβ plaque count was significantly reduced in Gluc/Fruc LP-fed mice and trended toward significance in Gluc LP-fed mice compared to their control counterparts (**Fig. 7B**). In contrast, total plaque area was significantly reduced in both Gluc/Fruc LP and Gluc LP compared to Con (**Fig. 7C**). There were no differences in microglia or astrocyte populations in the cortex, nor was there a difference in the interaction between those cells and Aβ plaques (**Figs. S7B-E**). To determine if reductions in plaques were due to a change in autophagy, we measured the expression of autophagy-related proteins in the cortex (**Figs. S8A-F**). Although autophagy was reduced overall compared to 4-month old controls (dotted lines in **Fig. S8B-F**), there was no effect of diet on autophagy. This suggests that reductions in plaques are likely due to a reduction in plaque formation as opposed to an increase in plaque clearance.

Ultimately, these results show that although the metabolic response of APP/PS1 females to dietary changes is blunted as compared to C57BL/6J mice, their metabolic health is still responsive to protein restriction and carbohydrate quality. Further, dietary protein impacts AD pathology, with LP-fed mice showing smaller and fewer plaques. Interestingly there was a divergence between this reduction in AD pathology and spatial memory impairment, with spatial memory differences between groups being driven by carbohydrate quality.

### Low protein diet improves metabolic health in male mice universally, but the female response to LP diets is defined by carbohydrate type

To determine how these diets affected health overall, we calculated composite z-scores for different categories of metabolic health C57BL/6J mice and AD pathology in APP/PS1 mice. **Table S5** lists which measures were included in calculating the composite z-score for each graph. Body composition was impacted primarily by carbohydrates in female mice, with control diets containing glucose monosaccharides having the most detrimental effects (**Fig. 8A**). In contrast, male mice had improved body composition in both Fruc Con and LP-fed mice. Thermogenesis was unaltered by carbohydrate type in the control context in both female and male mice, but there was an overall increase in thermogenesis in LP-fed mice, except for Fruc LP females and Gluc LP males (**Fig. 8B**). Glucose homeostasis was solely affected by protein, with improved glucose homeostasis in both female and male mice fed LP diets (**Fig. 8C**). In contrast, lipid homeostasis was primarily impacted by carbohydrate type, with fructose monosaccharides worsening lipid homeostasis in both female and male mice, particularly in the LP context (**Fig. 8D**). Female mice also exhibited effects of protein intake with LP diet improving lipid homeostasis in sucrose-fed mice with the opposite effect in fructose-fed mice. Taking into consideration all phenotypic measures of metabolic health, we found that protein intake had the strongest impact, which varied depending on carbohydrate content but had the strongest improvement on health in glucose/fructose and glucose-fed female and male mice (**Fig. 8E**). In APP/PS1 mice, there was no overall effect of carbohydrate or protein on AD pathology despite differences found in individual AD metrics (**Fig. 8F**).

**Figure 8:**
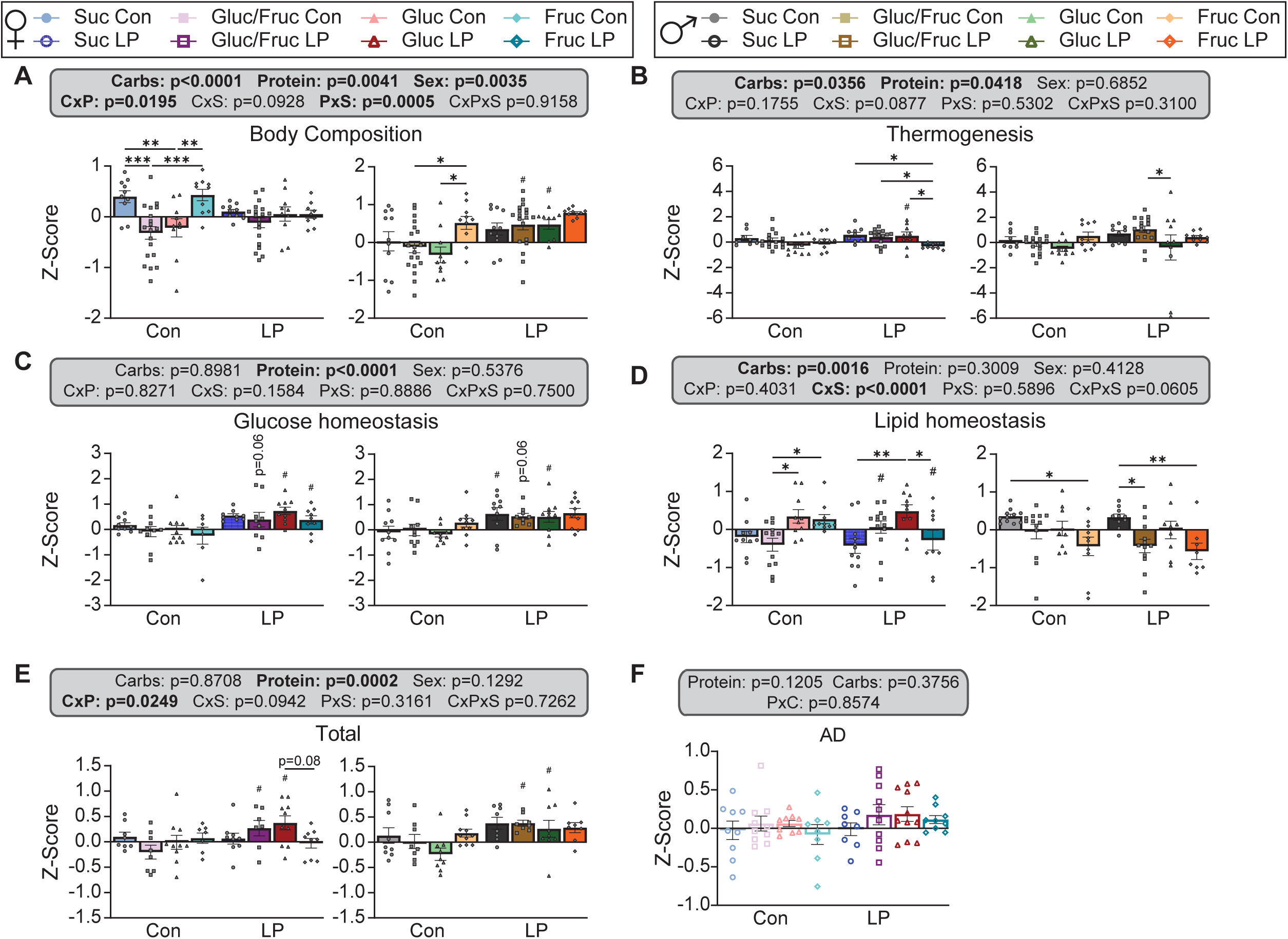
Composite z-scores of metabolic health. A-E) Composite z-scores of metabolic health: (A) body composition, (B) thermogenesis, (C) glucose homeostasis, (D) lipid homeostasis, and (E) overall. F) Composite z-score of AD phenotyping in APP/PS1 mice. n=7-19. All comparisons in this figure: asterisks denote differences between carbohydrate types within control or LP diets with *p<0.05, **p<0.01, ***p<0.001, ****p<0.0001, Tukey test post 2-way ANOVA. Hashes denote significant differences (p<0.05, Tukey test post 2-way ANOVA) in LP diets compared to their corresponding control counterparts. Exact p-values are provided in Table S3.

## Discussion

Low protein (LP) diets have been shown to robustly improve metabolic health in mice ^24-33,50^. However, most studies to date have utilized a standardized diet, with corn as the primary carbohydrate source. While this standardization strengthens reproducibility, humans routinely consume a much broader range of carbohydrates. Thus, effective translation of LP diets or even dietary recommendations requires that we understand how dietary carbohydrate quality impacts both the efficacy of LP diets and the ultimate effects of these diets on health. We therefore investigated how four different carbohydrate compositions - sucrose, glucose, fructose, and a glucose/fructose mixture – impact health as well as the response to LP diets.

We found that LP diets generally improve metabolic health with regards to body composition, thermogenesis, glucose homeostasis, and lipid homeostasis in both male and female C57BL/6J mice. Consistent with previous work, we observed that male mice have more robust metabolic benefits from consumption of an LP diet. However, we observed that carbohydrate quality also impacted metabolic health, especially in female mice. As we expected based on many studies, including Wali *et al.* ^50^, we observed that consumption of a 1:1 mixture of glucose and fructose, which is comparable to a diet high in high-fructose corn syrup, often resulted in worse health outcomes. As we hypothesized, we found that dietary carbohydrate quality does impact the response to LP diets.

Previous work in LP diets have found that while male mice respond strongly to PR, exhibiting reduced body weight and adiposity, female mice are more resistant to improvements to body composition ^24,25,27,28,34,61,64^. This occurs despite mice on LP diets having equal or greater caloric intake, and is likely due to increased Fgf21 expression, iWAT being, and energy expenditure ^24,28,34,61,64^. Here, we find that LP diets are effective at improving body composition and thermogenesis in male mice, but the effects of an LP diet on female mice vary with carbohydrate type. Protein intake correlated positively with body mass, lean mass, fat mass, and adiposity in male mice across all four carbohydrate types, resulting in reductions in LP-fed mice, but this was not the case in females. In female mice, protein intake was positively correlated with fat mass and adiposity only in the glucose and 1:1 glucose/fructose diets, whereas the opposite relationship was seen in diets containing sucrose or fructose. Interestingly, carbohydrate type had its own independent effects as well; fructose-only diets improved body weight and thermogenesis in both female and male mice in both control and LP contexts, while the 1:1 glucose/fructose diet worsened body weight in female mice. Unlike previous work, we did not see significantly worsened body composition in the diet with the highest level of fructose as in studies with fructose supplementation ^65,66^; instead these results were more similar to our 1:1 glucose/fructose diet group.

In addition to improvements in body composition and thermogenesis, LP diets have been shown to robustly improve glucose homeostasis ^24,25,34,64^. Here, this effect was prevalent in both sexes, with all LP-fed mice having improved glucose tolerance regardless of carbohydrate type, and some groups exhibiting improvements in insulin sensitivity. We also show that glucose metabolism is altered by carbohydrate type, with diets containing only glucose or only fructose tending to have improved glucose tolerance and insulin sensitivity. However, unlike body composition, we do not see any detrimental effects of 1:1 glucose/fructose diet on glucose homeostasis. Previous work on fructose supplementation found that increased fructose intake is accompanied by worsened glucose tolerance and insulin sensitivity ^40,66-69^, which we do not see here. One interesting finding is that of a “healthy” fatty liver in LP-fed mice. Mice of LP have increased lipid droplet and triglyceride levels in their livers, regardless of carbohydrate type, which would ordinarily be presumed to be negative for metabolic health. However, not only do these mice exhibit improved glucose tolerance (which is not ordinarily seen in fatty liver disease), plasma levels of AST and ALT are either unchanged or decreased. Interestingly, protein intake had minor impacts on other aspects of lipid homeostasis while carbohydrate type had robust, sex-dependent effects. Previous literature has shown increased hepatic TG levels in fructose supplementation ^65,66,68^, which is also true in our male mice compared to sucrose-fed mice. Softic *et al.* also found increases in hepatic fatty acid synthesis in fructose-fed mice ^40^, which we see strongly in our fructose-only females and males. In both females and males, diets containing fructose monosaccharides increased triglyceride release as determined via a P-407 assay.

LP diets or the restriction of the branched-chain amino acids improves cognition and AD pathology in mouse models ^30,31,51-53^. Our AD model showed blunted metabolic effects of both carbohydrates and protein, with there being overall benefits of LP diets while Gluc/Fruc diet worsened many aspects of metabolic health. Despite this, Aβ burden was ameliorated by LP as in previous studies. This was especially prominent in diets containing glucose. Unlike previous work, we did not see any differences in microglia or astrocyte populations or autophagy despite the decrease in amyloid burden, suggesting that plaque reduction may be driven by decreased amyloid production rather than increased clearance. Finally, we find worsened spatial memory in Gluc/Fruc fed mice, which is consistent with our findings that Gluc/Fruc mice tend to have worsened metabolic health and prior studies showing that fructose supplementation worsens AD pathology and cognition ^47,48^.

A consistent difference between our results and previous work on high fructose consumption is that we find our fructose-only diets to be beneficial for some aspects of metabolic health in both female and male mice. We hypothesize that the benefits of the fructose-only diet may be due to incomplete absorption of fructose. Prior work in humans has found that high fructose consumption in the absence of other sugars may result in incomplete absorption of the fructose ^70^. If incomplete fructose absorption is present in the current study, that may explain how fructose-only-fed mice are able to consume more calories and still lose body weight and adiposity, without a significant increase in energy expenditure. Future work should utilize bomb calorimetry to determine if this is true in the current dietary paradigm.

These results have important translational implications. Previous work in humans has shown detrimental effects on both metabolic health and AD by high fructose diets ^35,38,39,42,45,46,71-77^. One study even found a greater response to fructose in women compared to men, which is consistent with our findings that female mice are more responsive to carbohydrate type than males ^73^. Regarding protein intake, human studies are more inconsistent than in animal models, with some studies showing benefits in metabolic health, particularly body weight and adiposity, with high protein intake ^10-13,58^ while others find improvements to metabolic health and reduced risk of age-related diseases with low protein intake ^20,21,78^. Thus, it isn’t unreasonable to assume that inconsistencies between clinical studies—such as study demographics or other dietary components—may be to blame for inconsistent results. Here, we show that carbohydrate context can modulate health as well as the response to LP diet, suggesting that LP diet paradigms in humans should also take carbohydrate, and likely fat, intake into consideration.

One limitation of this study is that only young C57BL/6J mice were used. While LP and fructose-only diets showed benefits here, we do not know if these benefits would be maintained over a longer period or in older mice. As we have previously shown that strain contributes to the metabolic response to LP diet ^24^ and others have shown variable responses to fructose in different strains ^79-82^, it is important for future studies to test the robustness of our results in different strains or heterogeneous mice to determine if the benefits of LP diets are retained in strains that may be more sensitive to high-fructose diets. Finally, behavioral assay tests were underpowered, which limits ability to detect significance. Given the current results, it is possible that more robust changes to cognition may be detectable with a larger n. Finally, there are limitations in the translatability of the improvements we saw in fructose-only diets, as a fructose-only diet in humans would be impractical and hard to adhere to. There are no foods that contain only fructose, although certain foods such as fruits are proportionally much higher in fructose than others. As such, a fructose-only diet would have to be manufactured, likely making it a quite expensive diet and difficult to adhere to, thus eliminating the primary advantages of LP diets when compared to caloric restriction or pharmaceutical interventions. Thus, the translational potential lies instead in the novel findings regarding the ability of carbohydrates to modulate response to LP diet in females.

In conclusion, LP diets were effective regardless of carbohydrate context in male C57BL/6J mice. In female mice, however, carbohydrate type defined the response to LP diet; in the current study, female C57BL/6J mice were more responsive to LP diet than in previous studies under certain carbohydrate contexts, particularly with regards to body composition. This has important translational implications, as humans who consume an LP diet may be replacing calories from protein with carbohydrates from a variety of sources, which may contribute to the variability of reported responses to dietary protein intake in clinical trials.

## DATA AVAILABILITY STATEMENT

The data that support the plots within this article and other findings of this study, including full scans of western blot images, are provided as Source Data files.

## ACKNOWLEDGEMENTS

We would like to thank all members of the Lamming lab for their assistance and input. The Lamming laboratory is supported in part by the NIH/NIA (AG056771, AG081482, AG084156, AG085898 and AG094153 to D.W.L.), NIH/NIDDK (DK125859 to D.W.L.) and startup funds from the University of Wisconsin-Madison School of Medicine and Public Health and Department of Medicine to D.W.L. C-Y.Y. was supported in part by a NIA K99/R00 award (K99AG084921). R.B. was supported by F31AG081115. RB is a UW Distinguished Research Fellow; support for this research was provided by the University of Wisconsin-Madison Office of the Vice Chancellor for Research with funding from the Wisconsin Alumni Research Foundation. The authors used the UW Carbone Cancer Center Experimental Animal Pathology Laboratory (P30CA014520) for tissue sectioning and histology. D.W.L. is a member of the Wisconsin Nathan Shock Center of Excellence in the Basic Biology of Aging, P30 AG092586. The Lamming lab was supported in part by the U.S. Department of Veterans Affairs (IS1-BX005524), and this work was supported using facilities and resources from the William S. Middleton Memorial Veterans Hospital. The content is solely the responsibility of the authors and does not necessarily represent the official views of the NIH. This work does not represent the views of the Department of Veterans Affairs or the United States Government.

## References

1. Chew, N.W.S., Ng, C.H., Tan, D.J.H., Kong, G., Lin, C., Chin, Y.H., Lim, W.H., Huang, D.Q., Quek, J., Fu, C.E., et al. (2023). The global burden of metabolic disease: Data from 2000 to 2019. Cell Metab 35, 414–428 e413. 10.1016/j.cmet.2023.02.003..

2. Zhang, H., Zhou, X.D., Shapiro, M.D., Lip, G.Y.H., Tilg, H., Valenti, L., Somers, V.K., Byrne, C.D., Targher, G., Yang, W., et al. (2024). Global burden of metabolic diseases, 1990-2021. Metabolism 160, 155999. 10.1016/j.metabol.2024.155999.

3. Abubakar, M., Giri, A., Goel, F., Khan, M., Gupta, J., Kumar, D., Kaushik, M., Rai, S.N., and Kumar, N. (2025). Diabetes, Alzheimer’s Disease Risk Factors, and the Cafeteria Diet: A Comprehensive Review. Curr Neuropharmacol. 10.2174/011570159X384737250626094315.

4. Agrawal, R., Reno, C.M., Sharma, S., Christensen, C., Huang, Y., and Fisher, S.J. (2021). Insulin action in the brain regulates both central and peripheral functions. Am J Physiol Endocrinol Metab 321, E156–E163. 10.1152/ajpendo.00642.2020.

5. Ahtiluoto, S., Polvikoski, T., Peltonen, M., Solomon, A., Tuomilehto, J., Winblad, B., Sulkava, R., and Kivipelto, M. (2010). Diabetes, Alzheimer disease, and vascular dementia: a population-based neuropathologic study. Neurology 75, 1195–1202. 10.1212/WNL.0b013e3181f4d7f8.

6. Darst, B.F., Lu, Q., Johnson, S.C., and Engelman, C.D. (2019). Integrated analysis of genomics, longitudinal metabolomics, and Alzheimer’s risk factors among 1,111 cohort participants. Genet Epidemiol 43, 657–674. 10.1002/gepi.22211.

7. Gonzalez, A., Calfio, C., Churruca, M., and Maccioni, R.B. (2022). Glucose metabolism and AD: evidence for a potential diabetes type 3. Alzheimers Res Ther 14, 56. 10.1186/s13195-022-00996-8.

8. Hamze, R., Delangre, E., Tolu, S., Moreau, M., Janel, N., Bailbe, D., and Movassat, J. (2022). Type 2 Diabetes Mellitus and Alzheimer’s Disease: Shared Molecular Mechanisms and Potential Common Therapeutic Targets. Int J Mol Sci 23. 10.3390/ijms232315287.

9. Sims-Robinson, C., Kim, B., Rosko, A., and Feldman, E.L. (2010). How does diabetes accelerate Alzheimer disease pathology? Nat Rev Neurol 6, 551-559. 10.1038/nrneurol.2010.130.

10. Due, A., Toubro, S., Skov, A.R., and Astrup, A. (2004). Effect of normal-fat diets, either medium or high in protein, on body weight in overweight subjects: a randomised 1-year trial. Int J Obes Relat Metab Disord 28, 1283–1290. 10.1038/sj.ijo.0802767.

11. Skov, A.R., Toubro, S., Ronn, B., Holm, L., and Astrup, A. (1999). Randomized trial on protein vs carbohydrate in ad libitum fat reduced diet for the treatment of obesity. Int J Obes Relat Metab Disord 23, 528–536. 10.1038/sj.ijo.0800867.

12. Weigle, D.S., Breen, P.A., Matthys, C.C., Callahan, H.S., Meeuws, K.E., Burden, V.R., and Purnell, J.Q. (2005). A high-protein diet induces sustained reductions in appetite, ad libitum caloric intake, and body weight despite compensatory changes in diurnal plasma leptin and ghrelin concentrations. Am J Clin Nutr 82, 41–48. 10.1093/ajcn.82.1.41.

13. Campos-Nonato, I., Hernandez, L., and Barquera, S. (2017). Effect of a High-Protein Diet versus Standard-Protein Diet on Weight Loss and Biomarkers of Metabolic Syndrome: A Randomized Clinical Trial. Obes Facts 10, 238–251. 10.1159/000471485.

14. Larsen, T.M., Dalskov, S.M., van Baak, M., Jebb, S.A., Papadaki, A., Pfeiffer, A.F., Martinez, J.A., Handjieva-Darlenska, T., Kunesova, M., Pihlsgard, M., et al. (2010). Diets with high or low protein content and glycemic index for weight-loss maintenance. N Engl J Med 363, 2102-2113. 10.1056/NEJMoa1007137.

15. Levine, M.E., Suarez, J.A., Brandhorst, S., Balasubramanian, P., Cheng, C.W., Madia, F., Fontana, L., Mirisola, M.G., Guevara-Aguirre, J., Wan, J., et al. (2014). Low protein intake is associated with a major reduction in IGF-1, cancer, and overall mortality in the 65 and younger but not older population. Cell Metab 19, 407-417. 10.1016/j.cmet.2014.02.006.

16. Lagiou, P., Sandin, S., Weiderpass, E., Lagiou, A., Mucci, L., Trichopoulos, D., and Adami, H.O. (2007). Low carbohydrate-high protein diet and mortality in a cohort of Swedish women. J Intern Med 261, 366-374. 10.1111/j.1365-2796.2007.01774.x.

17. Linn, T., Santosa, B., Gronemeyer, D., Aygen, S., Scholz, N., Busch, M., and Bretzel, R.G. (2000). Effect of long-term dietary protein intake on glucose metabolism in humans. Diabetologia 43, 1257–1265. 10.1007/s001250051521.

18. Sluijs, I., Beulens, J.W., van der, A.D., Spijkerman, A.M., Grobbee, D.E., and van der Schouw, Y.T. (2010). Dietary intake of total, animal, and vegetable protein and risk of type 2 diabetes in the European Prospective Investigation into Cancer and Nutrition (EPIC)-NL study. Diabetes Care 33, 43–48. 10.2337/dc09-1321.

19. van Nielen, M., Feskens, E.J., Mensink, M., Sluijs, I., Molina, E., Amiano, P., Ardanaz, E., Balkau, B., Beulens, J.W., Boeing, H., et al. (2014). Dietary protein intake and incidence of type 2 diabetes in Europe: the EPIC-InterAct Case-Cohort Study. Diabetes Care 37, 1854–1862. 10.2337/dc13-2627.

20. Nicolaisen, T.S., Lyster, A.E., Sjoberg, K.A., Haas, D.T., Voldstedlund, C.T., Lundsgaard, A.M., Jensen, J.K., Madsen, E.M., Nielsen, C.K., Bloch-Ibenfeldt, M., et al. (2025). Dietary protein restriction elevates FGF21 levels and energy requirements to maintain body weight in lean men. Nat Metab 7, 602–616. 10.1038/s42255-025-01236-7.

21. Ferraz-Bannitz, R., Beraldo, R.A., Peluso, A.A., Dall, M., Babaei, P., Foglietti, R.C., Martins, L.M., Gomes, P.M., Marchini, J.S., Suen, V.M.M., et al. (2022). Dietary Protein Restriction Improves Metabolic Dysfunction in Patients with Metabolic Syndrome in a Randomized, Controlled Trial. Nutrients 14. 10.3390/nu14132670.

22. Fontana, L., Cummings, N.E., Arriola Apelo, S.I., Neuman, J.C., Kasza, I., Schmidt, B.A., Cava, E., Spelta, F., Tosti, V., Syed, F.A., et al. (2016). Decreased Consumption of Branched-Chain Amino Acids Improves Metabolic Health. Cell Rep 16, 520-530. 10.1016/j.celrep.2016.05.092.

23. Lyster, A.E., Frederiksen, A.S., Jensen, J.K., Lundsgaard, A., Andersen, N.R., Jorgensen, K.S., Weber, D., Nachtigall, L., Richter, E.A., Fritzen, A.M., and Kiens, B. (2026). Dietary Protein Reduction During Isocaloric Conditions Reduces Body Weight in Men With Overweight or Obesity. Obesity (Silver Spring). 10.1002/oby.70248.

24. Green, C.L., Pak, H.H., Richardson, N.E., Flores, V., Yu, D., Tomasiewicz, J.L., Dumas, S.N., Kredell, K., Fan, J.W., Kirsh, C., et al. (2022). Sex and genetic background define the metabolic, physiologic, and molecular response to protein restriction. Cell Metab 34, 209–226 e205. 10.1016/j.cmet.2021.12.018.

25. Hill, C.M., Albarado, D.C., Coco, L.G., Spann, R.A., Khan, M.S., Qualls-Creekmore, E., Burk, D.H., Burke, S.J., Collier, J.J., Yu, S., et al. (2022). FGF21 is required for protein restriction to extend lifespan and improve metabolic health in male mice. Nat Commun 13, 1897. 10.1038/s41467-022-29499-8.

26. Lamming, D.W., Cummings, N.E., Rastelli, A.L., Gao, F., Cava, E., Bertozzi, B., Spelta, F., Pili, R., and Fontana, L. (2015). Restriction of dietary protein decreases mTORC1 in tumors and somatic tissues of a tumor-bearing mouse xenograft model. Oncotarget 6, 31233-31240. 10.18632/oncotarget.5180.

27. Solon-Biet, S.M., Mitchell, S.J., Coogan, S.C., Cogger, V.C., Gokarn, R., McMahon, A.C., Raubenheimer, D., de Cabo, R., Simpson, S.J., and Le Couteur, D.G. (2015). Dietary Protein to Carbohydrate Ratio and Caloric Restriction: Comparing Metabolic Outcomes in Mice. Cell Rep 11, 1529–1534. 10.1016/j.celrep.2015.05.007.

28. Laeger, T., Henagan, T.M., Albarado, D.C., Redman, L.M., Bray, G.A., Noland, R.C., Munzberg, H., Hutson, S.M., Gettys, T.W., Schwartz, M.W., and Morrison, C.D. (2014). FGF21 is an endocrine signal of protein restriction. J Clin Invest 124, 3913–3922. 10.1172/JCI74915.

29. Maida, A., Chan, J.S.K., Sjoberg, K.A., Zota, A., Schmoll, D., Kiens, B., Herzig, S., and Rose, A.J. (2017). Repletion of branched chain amino acids reverses mTORC1 signaling but not improved metabolism during dietary protein dilution. Mol Metab 6, 873-881. 10.1016/j.molmet.2017.06.009.

30. Babygirija, R., Sonsalla, M.M., Mill, J., James, I., Han, J.H., Green, C.L., Calubag, M.F., Wade, G., Tobon, A., Michael, J., et al. (2024). Protein restriction slows the development and progression of pathology in a mouse model of Alzheimer’s disease. Nat Commun 15, 5217. 10.1038/s41467-024-49589-z.

31. Parrella, E., Maxim, T., Maialetti, F., Zhang, L., Wan, J., Wei, M., Cohen, P., Fontana, L., and Longo, V.D. (2013). Protein restriction cycles reduce IGF-1 and phosphorylated Tau, and improve behavioral performance in an Alzheimer’s disease mouse model. Aging Cell 12, 257–268. 10.1111/acel.12049.

32. Richardson, N.E., Konon, E.N., Schuster, H.S., Mitchell, A.T., Boyle, C., Rodgers, A.C., Finke, M., Haider, L.R., Yu, D., Flores, V., et al. (2021). Lifelong restriction of dietary branched-chain amino acids has sex-specific benefits for frailty and lifespan in mice. Nat Aging 1, 73–86. 10.1038/s43587-020-00006-2.

33. Solon-Biet, S.M., McMahon, A.C., Ballard, J.W., Ruohonen, K., Wu, L.E., Cogger, V.C., Warren, A., Huang, X., Pichaud, N., Melvin, R.G., et al. (2014). The ratio of macronutrients, not caloric intake, dictates cardiometabolic health, aging, and longevity in ad libitum-fed mice. Cell Metab 19, 418-430. 10.1016/j.cmet.2014.02.009.

34. Knopf, B.A., Grunow, I., Anderson, B., Rihawi, T., Sonsalla, M.M., Calubag, M.F., Babygirija, R., Liu, Y., Xiao, F., Yeh, C.Y., and Lamming, D.W. (2026). Female resistance to the metabolic benefits of protein restriction is reversed by ovariectomy in mice. bioRxiv. 10.64898/2026.03.31.715667.

35. Tappy, L. (2018). Fructose-containing caloric sweeteners as a cause of obesity and metabolic disorders. J Exp Biol 221. 10.1242/jeb.164202.

36. Clemente-Suarez, V.J., Beltran-Velasco, A.I., Redondo-Florez, L., Martin-Rodriguez, A., and Tornero-Aguilera, J.F. (2023). Global Impacts of Western Diet and Its Effects on Metabolism and Health: A Narrative Review. Nutrients 15. 10.3390/nu15122749.

37. Demaria, T.M., Crepaldi, L.D., Costa-Bartuli, E., Branco, J.R., Zancan, P., and Sola-Penna, M. (2023). Once a week consumption of Western diet over twelve weeks promotes sustained insulin resistance and non-alcoholic fat liver disease in C57BL/6 J mice. Sci Rep 13, 3058. 10.1038/s41598-023-30254-2.

38. Tappy, L., and Rosset, R. (2019). Health outcomes of a high fructose intake: the importance of physical activity. J Physiol 597, 3561-3571. 10.1113/JP278246.

39. Geidl-Flueck, B., Hochuli, M., Nemeth, A., Eberl, A., Derron, N., Kofeler, H.C., Tappy, L., Berneis, K., Spinas, G.A., and Gerber, P.A. (2021). Fructose- and sucrose-but not glucose-sweetened beverages promote hepatic de novo lipogenesis: A randomized controlled trial. J Hepatol 75, 46–54. 10.1016/j.jhep.2021.02.027.

40. Softic, S., Gupta, M.K., Wang, G.X., Fujisaka, S., O’Neill, B.T., Rao, T.N., Willoughby, J., Harbison, C., Fitzgerald, K., Ilkayeva, O., et al. (2017). Divergent effects of glucose and fructose on hepatic lipogenesis and insulin signaling. J Clin Invest 127, 4059–4074. 10.1172/JCI94585.

41. Fisher, F.M., Kim, M., Doridot, L., Cunniff, J.C., Parker, T.S., Levine, D.M., Hellerstein, M.K., Hudgins, L.C., Maratos-Flier, E., and Herman, M.A. (2017). A critical role for ChREBP-mediated FGF21 secretion in hepatic fructose metabolism. Mol Metab 6, 14-21. 10.1016/j.molmet.2016.11.008.

42. Herman, M.A., and Birnbaum, M.J. (2021). Molecular aspects of fructose metabolism and metabolic disease. Cell Metab 33, 2329–2354. 10.1016/j.cmet.2021.09.010.

43. Hu, Y., Semova, I., Sun, X., Kang, H., Chahar, S., Hollenberg, A.N., Masson, D., Hirschey, M.D., Miao, J., and Biddinger, S.B. (2018). Fructose and glucose can regulate mammalian target of rapamycin complex 1 and lipogenic gene expression via distinct pathways. J Biol Chem 293, 2006-2014. 10.1074/jbc.M117.782557.

44. Seneff, S., Wainwright, G., and Mascitelli, L. (2011). Nutrition and Alzheimer’s disease: the detrimental role of a high carbohydrate diet. Eur J Intern Med 22, 134–140. 10.1016/j.ejim.2010.12.017.

45. Johnson, R.J., Gomez-Pinilla, F., Nagel, M., Nakagawa, T., Rodriguez-Iturbe, B., Sanchez-Lozada, L.G., Tolan, D.R., and Lanaspa, M.A. (2020). Cerebral Fructose Metabolism as a Potential Mechanism Driving Alzheimer’s Disease. Front Aging Neurosci 12, 560865. 10.3389/fnagi.2020.560865.

46. Yan, J., Zheng, K., Zhang, X., and Jiang, Y. (2023). Fructose Consumption is Associated with a Higher Risk of Dementia and Alzheimer’s Disease: A Prospective Cohort Study. J Prev Alzheimers Dis 10, 186–192. 10.14283/jpad.2023.7.

47. Ormazabal, P., Ordenes-Constenla, P., Gherardelli, C., Bastias-Perez, M., Brito-Valenzuela, J., Flores-Opazo, M., Inestrosa, N.C., and Cisternas, P. (2026). Fructose Intake Is Associated with Brain Metabolic Reprogramming and Exacerbation of Alzheimer-like Alterations in APP/PS1 Mice. Int J Mol Sci 27. 10.3390/ijms27094113.

48. Anderson, R.A., Qin, B., Canini, F., Poulet, L., and Roussel, A.M. (2013). Cinnamon counteracts the negative effects of a high fat/high fructose diet on behavior, brain insulin signaling and Alzheimer-associated changes. PLoS One 8, e83243. 10.1371/journal.pone.0083243.

49. Pase, M.P., Himali, J.J., Jacques, P.F., DeCarli, C., Satizabal, C.L., Aparicio, H., Vasan, R.S., Beiser, A.S., and Seshadri, S. (2017). Sugary beverage intake and preclinical Alzheimer’s disease in the community. Alzheimers Dement 13, 955–964. 10.1016/j.jalz.2017.01.024.

50. Wali, J.A., Milner, A.J., Luk, A.W.S., Pulpitel, T.J., Dodgson, T., Facey, H.J.W., Wahl, D., Kebede, M.A., Senior, A.M., Sullivan, M.A., et al. (2021). Impact of dietary carbohydrate type and protein-carbohydrate interaction on metabolic health. Nat Metab 3, 810–828. 10.1038/s42255-021-00393-9.

51. Babygirija, R., Green, C.L., Sonsalla, M.M., Le, I.M.F., Xiao, F., Yandell, S., Calubag, M.F., Trautman, M.E., Tobon, A., Matoska, R., et al. (2026). Restriction of Individual Branched-Chain Amino Acids has Distinct Effects on the Development and Progression of Alzheimer’s Disease in 3xTg Mice. Adv Sci (Weinh), e15220. 10.1002/advs.202515220.

52. Siddik, M.A.B., Mullins, C.A., Kramer, A., Shah, H., Gannaban, R.B., Zabet-Moghaddam, M., Huebinger, R.M., Hegde, V.K., MohanKumar, S.M.J., MohanKumar, P.S., and Shin, A.C. (2022). Branched-Chain Amino Acids Are Linked with Alzheimer’s Disease-Related Pathology and Cognitive Deficits. Cells 11. 10.3390/cells11213523.

53. Tournissac, M., Vandal, M., Tremblay, C., Bourassa, P., Vancassel, S., Emond, V., Gangloff, A., and Calon, F. (2018). Dietary intake of branched-chain amino acids in a mouse model of Alzheimer’s disease: Effects on survival, behavior, and neuropathology. Alzheimers Dement (N Y) 4, 677–687. 10.1016/j.trci.2018.10.005.

54. Bellantuono, I., de Cabo, R., Ehninger, D., Di Germanio, C., Lawrie, A., Miller, J., Mitchell, S.J., Navas-Enamorado, I., Potter, P.K., Tchkonia, T., et al. (2020). A toolbox for the longitudinal assessment of healthspan in aging mice. Nat Protoc 15, 540–574. 10.1038/s41596-019-0256-1.

55. Yu, D., Yang, S.E., Miller, B.R., Wisinski, J.A., Sherman, D.S., Brinkman, J.A., Tomasiewicz, J.L., Cummings, N.E., Kimple, M.E., Cryns, V.L., and Lamming, D.W. (2018). Short-term methionine deprivation improves metabolic health via sexually dimorphic, mTORC1-independent mechanisms. FASEB J 32, 3471-3482. 10.1096/fj.201701211R.

56. Cummings, N.E., Williams, E.M., Kasza, I., Konon, E.N., Schaid, M.D., Schmidt, B.A., Poudel, C., Sherman, D.S., Yu, D., Arriola Apelo, S.I., et al. (2018). Restoration of metabolic health by decreased consumption of branched-chain amino acids. J Physiol 596, 623-645. 10.1113/JP275075.

57. Yu, D., Richardson, N.E., Green, C.L., Spicer, A.B., Murphy, M.E., Flores, V., Jang, C., Kasza, I., Nikodemova, M., Wakai, M.H., et al. (2021). The adverse metabolic effects of branched-chain amino acids are mediated by isoleucine and valine. Cell Metab 33, 905–922 e906. 10.1016/j.cmet.2021.03.025.

58. Larson, K.R., Russo, K.A., Fang, Y., Mohajerani, N., Goodson, M.L., and Ryan, K.K. (2017). Sex Differences in the Hormonal and Metabolic Response to Dietary Protein Dilution. Endocrinology 158, 3477–3487. 10.1210/en.2017-00331.

59. Bruss, M.D., Khambatta, C.F., Ruby, M.A., Aggarwal, I., and Hellerstein, M.K. (2010). Calorie restriction increases fatty acid synthesis and whole body fat oxidation rates. Am J Physiol Endocrinol Metab 298, E108-116. 10.1152/ajpendo.00524.2009.

60. Pak, H.H., Haws, S.A., Green, C.L., Koller, M., Lavarias, M.T., Richardson, N.E., Yang, S.E., Dumas, S.N., Sonsalla, M., Bray, L., et al. (2021). Fasting drives the metabolic, molecular and geroprotective effects of a calorie-restricted diet in mice. Nat Metab 3, 1327–1341. 10.1038/s42255-021-00466-9.

61. Hill, C.M., Laeger, T., Albarado, D.C., McDougal, D.H., Berthoud, H.R., Munzberg, H., and Morrison, C.D. (2017). Low protein-induced increases in FGF21 drive UCP1-dependent metabolic but not thermoregulatory endpoints. Sci Rep 7, 8209. 10.1038/s41598-017-07498-w.

62. Paolella, L.M., Mukherjee, S., Tran, C.M., Bellaver, B., Hugo, M., Luongo, T.S., Shewale, S.V., Lu, W., Chellappa, K., and Baur, J.A. (2020). mTORC1 restrains adipocyte lipolysis to prevent systemic hyperlipidemia. Mol Metab 32, 136–147. 10.1016/j.molmet.2019.12.003.

63. Sonsalla, M.M., and Lamming, D.W. (2023). Geroprotective interventions in the 3xTg mouse model of Alzheimer’s disease. Geroscience 45, 1343–1381. 10.1007/s11357-023-00782-w.

64. Hill, C.M., Laeger, T., Dehner, M., Albarado, D.C., Clarke, B., Wanders, D., Burke, S.J., Collier, J.J., Qualls-Creekmore, E., Solon-Biet, S.M., et al. (2019). FGF21 Signals Protein Status to the Brain and Adaptively Regulates Food Choice and Metabolism. Cell reports 27, 2934–2947 e2933. 10.1016/j.celrep.2019.05.022.

65. Zhao, H., Tian, Y., Zuo, Y., Zhang, X., Gao, Y., Wang, P., Sun, L., Zhang, H., and Liang, H. (2022). Nicotinamide riboside ameliorates high-fructose-induced lipid metabolism disorder in mice via improving FGF21 resistance in the liver and white adipose tissue. Food Funct 13, 12400–12411. 10.1039/d2fo01934e.

66. Ahmed Mustafa, Z., Hamed Ali, R., Rostum Ali, D., Abdulkarimi, R., Abdulkareem, N.K., and Akbari, A. (2021). The combination of ginger powder and zinc supplement improves the fructose-induced metabolic syndrome in rats by modulating the hepatic expression of NF-kappaB, mTORC1, PPAR-alpha SREBP-1c, and Nrf2. J Food Biochem 45, e13546. 10.1111/jfbc.13546.

67. Baena, M., Sanguesa, G., Davalos, A., Latasa, M.J., Sala-Vila, A., Sanchez, R.M., Roglans, N., Laguna, J.C., and Alegret, M. (2016). Fructose, but not glucose, impairs insulin signaling in the three major insulin-sensitive tissues. Sci Rep 6, 26149. 10.1038/srep26149.

68. Wang, H., Sun, R.Q., Zeng, X.Y., Zhou, X., Li, S., Jo, E., Molero, J.C., and Ye, J.M. (2015). Restoration of autophagy alleviates hepatic ER stress and impaired insulin signalling transduction in high fructose-fed male mice. Endocrinology 156, 169-181. 10.1210/en.2014-1454.

69. Softic, S., Stanhope, K.L., Boucher, J., Divanovic, S., Lanaspa, M.A., Johnson, R.J., and Kahn, C.R. (2020). Fructose and hepatic insulin resistance. Crit Rev Clin Lab Sci 57, 308–322. 10.1080/10408363.2019.1711360.

70. Truswell, A.S., Seach, J.M., and Thorburn, A.W. (1988). Incomplete absorption of pure fructose in healthy subjects and the facilitating effect of glucose. Am J Clin Nutr 48, 1424–1430. 10.1093/ajcn/48.6.1424.

71. Jung, S., Bae, H., Song, W.S., and Jang, C. (2022). Dietary Fructose and Fructose-Induced Pathologies. Annu Rev Nutr 42, 45–66. 10.1146/annurev-nutr-062220-025831.

72. Lakhan, S.E., and Kirchgessner, A. (2013). The emerging role of dietary fructose in obesity and cognitive decline. Nutr J 12, 114. 10.1186/1475-2891-12-114.

73. Low, W.S., Cornfield, T., Charlton, C.A., Tomlinson, J.W., and Hodson, L. (2018). Sex Differences in Hepatic De Novo Lipogenesis with Acute Fructose Feeding. Nutrients 10. 10.3390/nu10091263.

74. Roeb, E., and Weiskirchen, R. (2021). Fructose and Non-Alcoholic Steatohepatitis. Front Pharmacol 12, 634344. 10.3389/fphar.2021.634344.

75. Stanhope, K.L., Schwarz, J.M., and Havel, P.J. (2013). Adverse metabolic effects of dietary fructose: results from the recent epidemiological, clinical, and mechanistic studies. Curr Opin Lipidol 24, 198–206. 10.1097/MOL.0b013e3283613bca.

76. Tappy, L. (2018). Fructose metabolism and noncommunicable diseases: recent findings and new research perspectives. Curr Opin Clin Nutr Metab Care 21, 214–222. 10.1097/MCO.0000000000000460.

77. Taskinen, M.R., Packard, C.J., and Boren, J. (2019). Dietary Fructose and the Metabolic Syndrome. Nutrients 11. 10.3390/nu11091987.

78. Fontana, L., Weiss, E.P., Villareal, D.T., Klein, S., and Holloszy, J.O. (2008). Long-term effects of calorie or protein restriction on serum IGF-1 and IGFBP-3 concentration in humans. Aging Cell 7, 681–687. 10.1111/j.1474-9726.2008.00417.x.

79. Ahn, I.S., Lang, J.M., Olson, C.A., Diamante, G., Zhang, G., Ying, Z., Byun, H.R., Cely, I., Ding, J., Cohn, P., et al. (2020). Host Genetic Background and Gut Microbiota Contribute to Differential Metabolic Responses to Fructose Consumption in Mice. J Nutr 150, 2716-2728. 10.1093/jn/nxaa239.

80. Ahn, I.S., Yoon, J., Diamante, G., Cohn, P., Jang, C., and Yang, X. (2021). Disparate Metabolomic Responses to Fructose Consumption between Different Mouse Strains and the Role of Gut Microbiota. Metabolites 11. 10.3390/metabo11060342.

81. Pinhas, A., Aviel, M., Koen, M., Gurgov, S., Acosta, V., Israel, M., Kakuriev, L., Guskova, E., Fuzailov, I., Touzani, K., et al. (2012). Strain differences in sucrose- and fructose-conditioned flavor preferences in mice. Physiol Behav 105, 451–459. 10.1016/j.physbeh.2011.09.010.

82. Zhang, G., Byun, H.R., Ying, Z., Blencowe, M., Zhao, Y., Hong, J., Shu, L., Chella Krishnan, K., Gomez-Pinilla, F., and Yang, X. (2020). Differential metabolic and multi-tissue transcriptomic responses to fructose consumption among genetically diverse mice. Biochim Biophys Acta Mol Basis Dis 1866, 165569. 10.1016/j.bbadis.2019.165569.

