## Supplemental Figures and Table Legends for "Dietary sugar type determines the response to protein restriction in females, but not males"

Figure S1

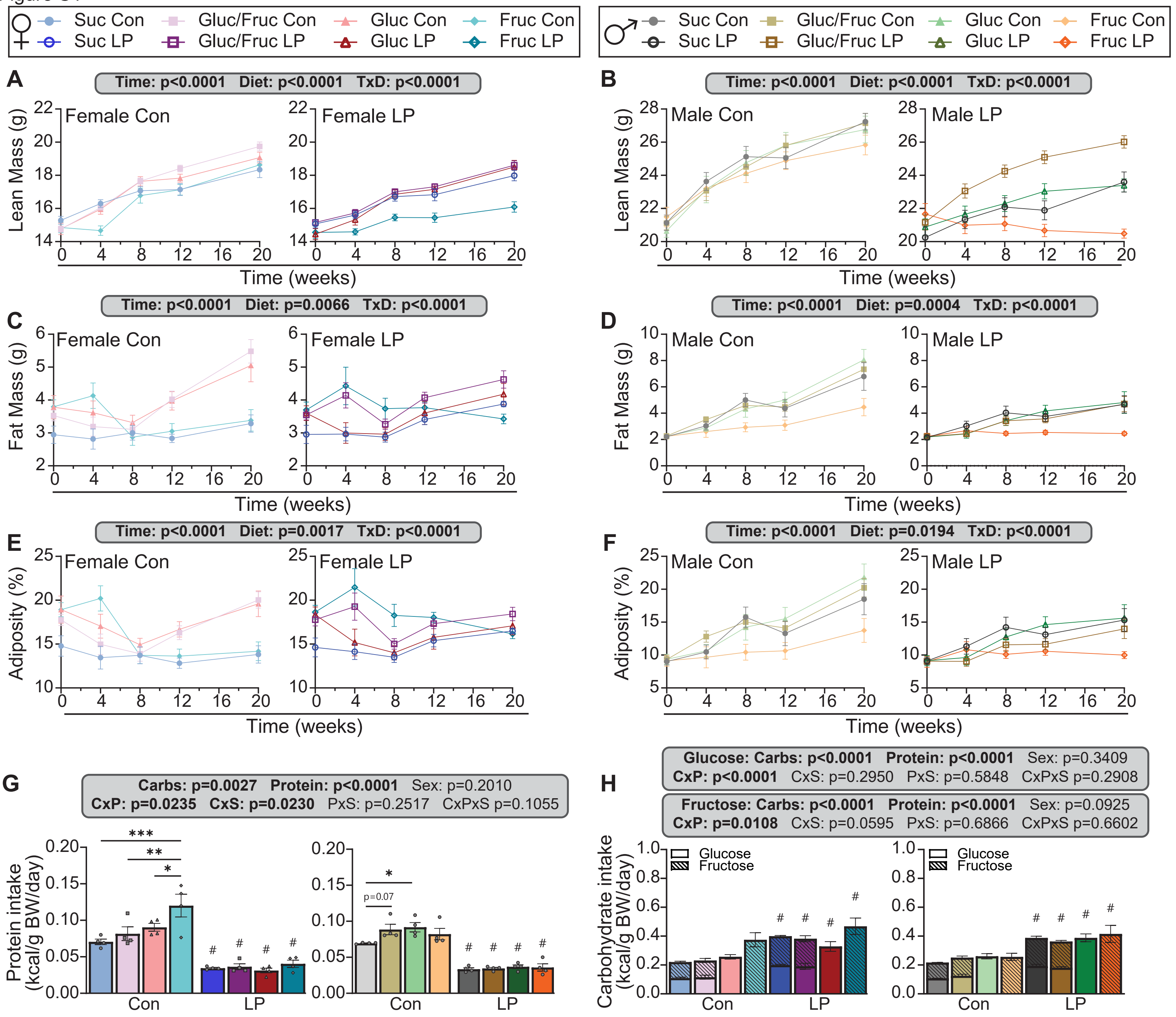
Suc Con
Suc LP
Gluc/Fruc Con
Gluc/Fruc LP
Gluc Con
Gluc LP
Fruc Con
Fruc LP
A
Time: p<0.0001 Diet: p<0.0001 TxD: p<0.0001
Female Con
Female LP
Lean Mass (g)
Time (weeks)
B
Time: p<0.0001 Diet: p<0.0001 TxD: p<0.0001
Male Con
Male LP
C
Time: p<0.0001 Diet: p=0.0066 TxD: p<0.0001
Fat Mass (g)
D
Time: p<0.0001 Diet: p=0.0004 TxD: p<0.0001
E
Time: p<0.0001 Diet: p=0.0017 TxD: p<0.0001
Adiposity (%)
F
Time: p<0.0001 Diet: p=0.0194 TxD: p<0.0001
G
Carbs: p=0.0027 Protein: p<0.0001 Sex: p=0.2010
CxP: p=0.0235 CxS: p=0.0230 PxS: p=0.2517 CxPxS p=0.1055
Protein intake (kcal/g BW/day)
p=0.07
Con
LP
H
Glucose: Carbs: p<0.0001 Protein: p<0.0001 Sex: p=0.3409
CxP: p<0.0001 CxS: p=0.2950 PxS: p=0.5848 CxPxS p=0.2908
Fructose: Carbs: p<0.0001 Protein: p<0.0001 Sex: p=0.0925
CxP: p=0.0108 CxS: p=0.0595 PxS: p=0.6866 CxPxS p=0.6602
Carbohydrate intake (kcal/g BW/day)
Glucose
Fructose

**Figure S1: Lean mass, fat mass, and adiposity accretion were blunted in fructose and LP-fed mice**.

(A-B) Lean mass over the course of the experiment in female (A) and male (B) mice.

(C-D) Fat mass over the course of the experiment in female (C) and male (D) mice.

(E-F) Adiposity over the course of the experiment in female (E) and male (F) mice.

(G) Protein intake normalized to body weight, calculated from home cage food consumption.

(H) Glucose (solid) and fructose (striped) consumption in female and male mice normalized to body weight, calculated from home cage food consumption.

Body weight and composition data n=10-19; home cage data n=4. All comparisons in this figure: asterisks denote differences between carbohydrate types within control or LP diets with *$p<0.05$, **$p<0.01$, ***$p<0.001$, ****$p<0.0001$, Tukey test post 2-way ANOVA. Hashes denote significant differences ($p<0.05$, Tukey test post 2-way ANOVA) in LP diets compared to their corresponding control counterparts. Exact p-values are provided in Table S3.

Supp Figure 2

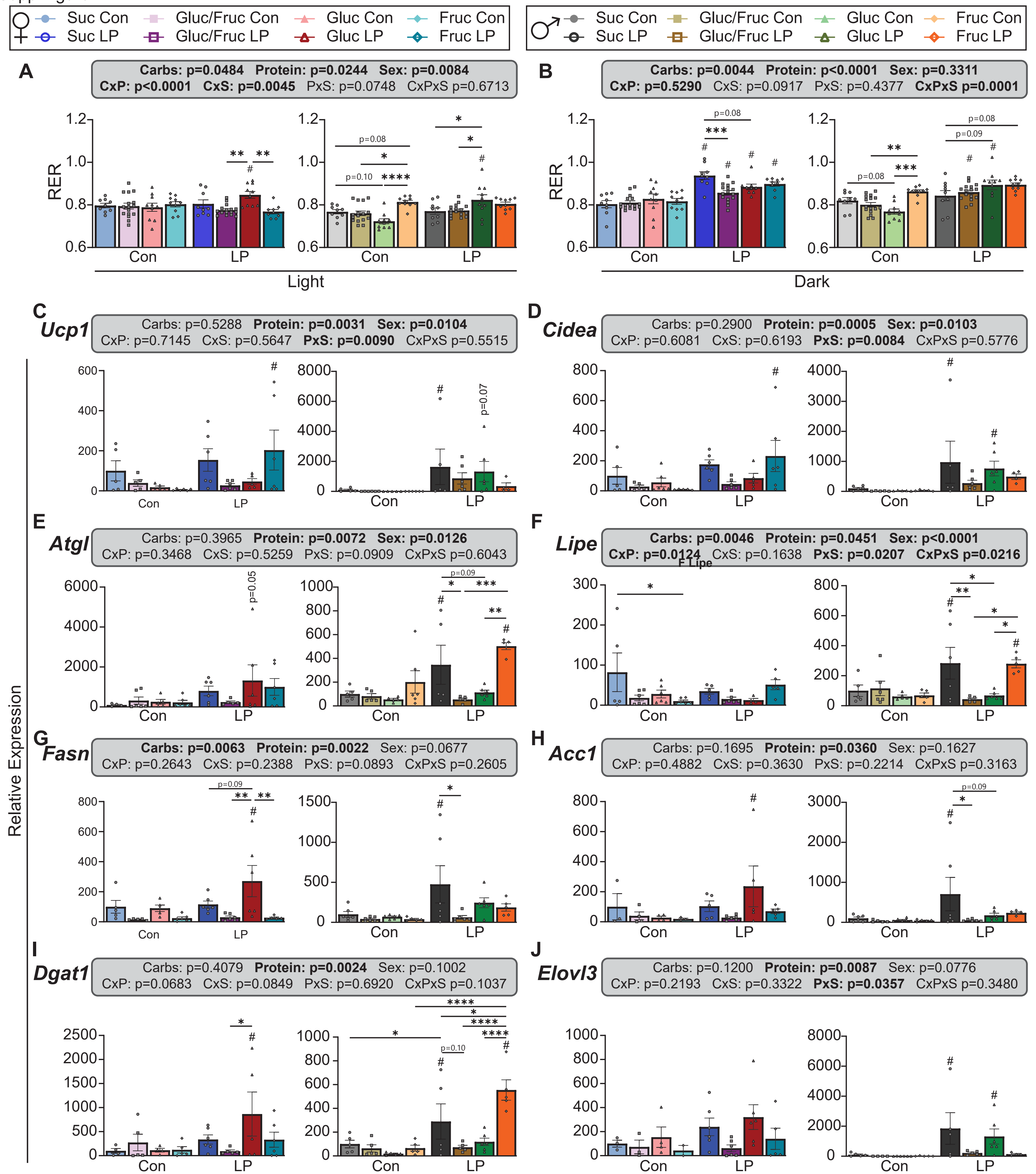

**Figure S2: Protein and carbohydrate intake impact thermogenic, lipogenic, and lipolytic gene expression in inguinal white adipose tissue.**

(A-B) Respiratory exchange ratio (RER) calculated from singly housed mice in Oxymax/CLAMS metabolic cages during the light (A) and dark (B) period. n=10-19

(C-J) Individual results for iWAT qPCRs in the heat map in Fig. 2G; genes measured include *Ucp1* (C), *Cidea* (D), *Atgl* (E), *Lipe* (F), *Fasn* (G), *Acc1* (H), *Dgat1* (I), and *Elovl3* (J). n=4-5

All comparisons in this figure: asterisks denote differences between carbohydrate types within control or LP diets with *p<0.05, **p<0.01, ***p<0.001, ****p<0.0001, Tukey test post 2-way ANOVA. Hashes denote significant differences (p<0.05, Tukey test post 2-way ANOVA) in LP diets compared to their corresponding control counterparts. Exact p-values are provided in Table S3.

Supp Figure 3

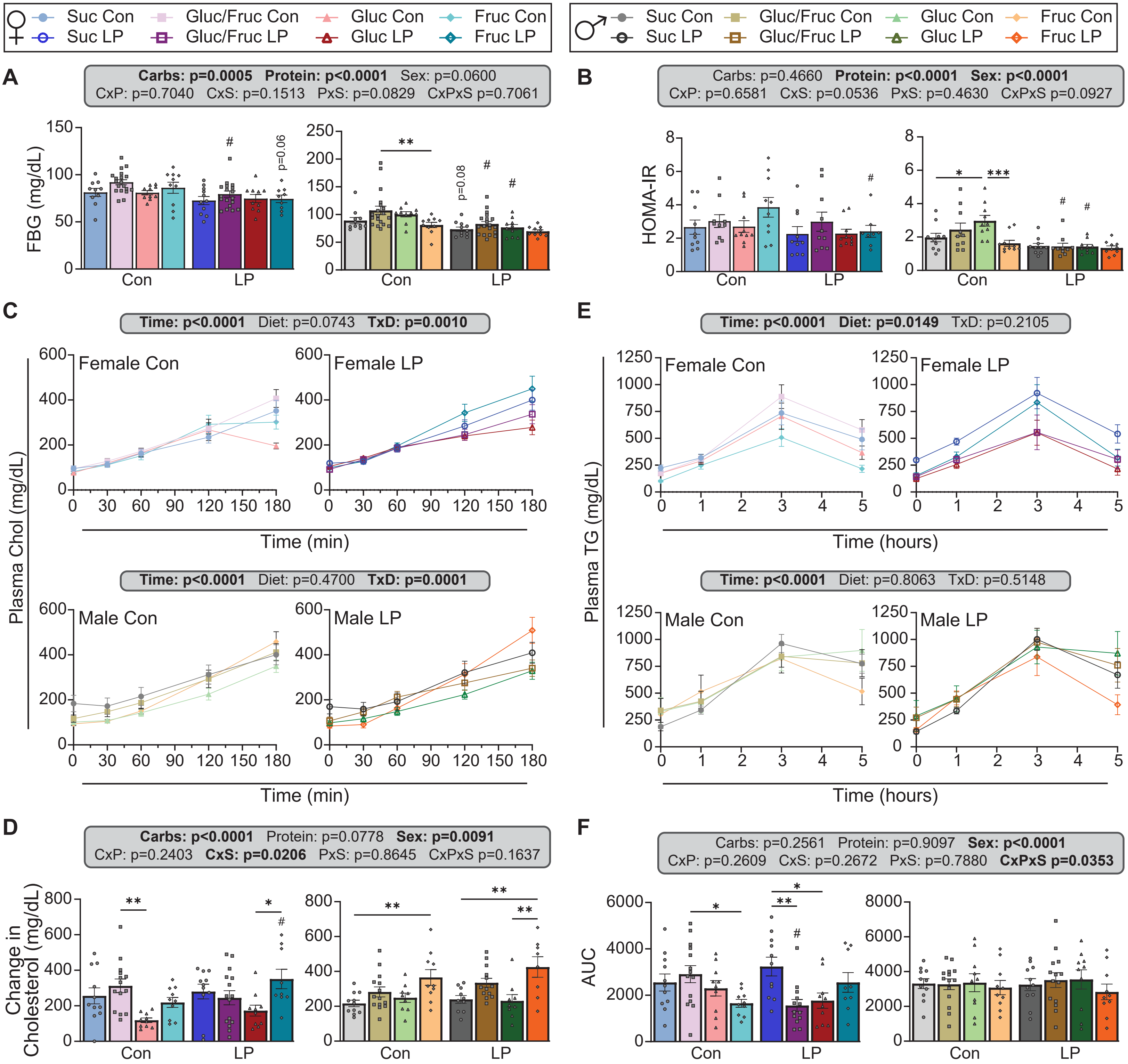

**Figure S3: Fasting blood glucose is primarily impacted by protein intake, but lipid homeostasis is affected by carbohydrate type.**

(A) Fasting blood glucose levels in female and male mice after a 16 hour fast.

(B) HOMA-IR calculation in female and male mice.

(C-D) P-407 assays were performed in female and male mice (C) as an *in vivo* approximation of lipid homeostasis and change in cholesterol over the 3 hour assay quantified (D).

(E-F) A lipid tolerance test was performed (E) and area under the curve measured (F) to further assess lipid homeostasis.

All comparisons in this figure: asterisks denote differences between carbohydrate types within control or LP diets with *$p<0.05$, **$p<0.01$, ***$p<0.001$, ****$p<0.0001$, Tukey test post 2-way ANOVA. Hashes denote significant differences ($p<0.05$, Tukey test post 2-way ANOVA) in LP diets compared to their corresponding control counterparts. Exact p-values are provided in Table S3.

Supp Figure 4

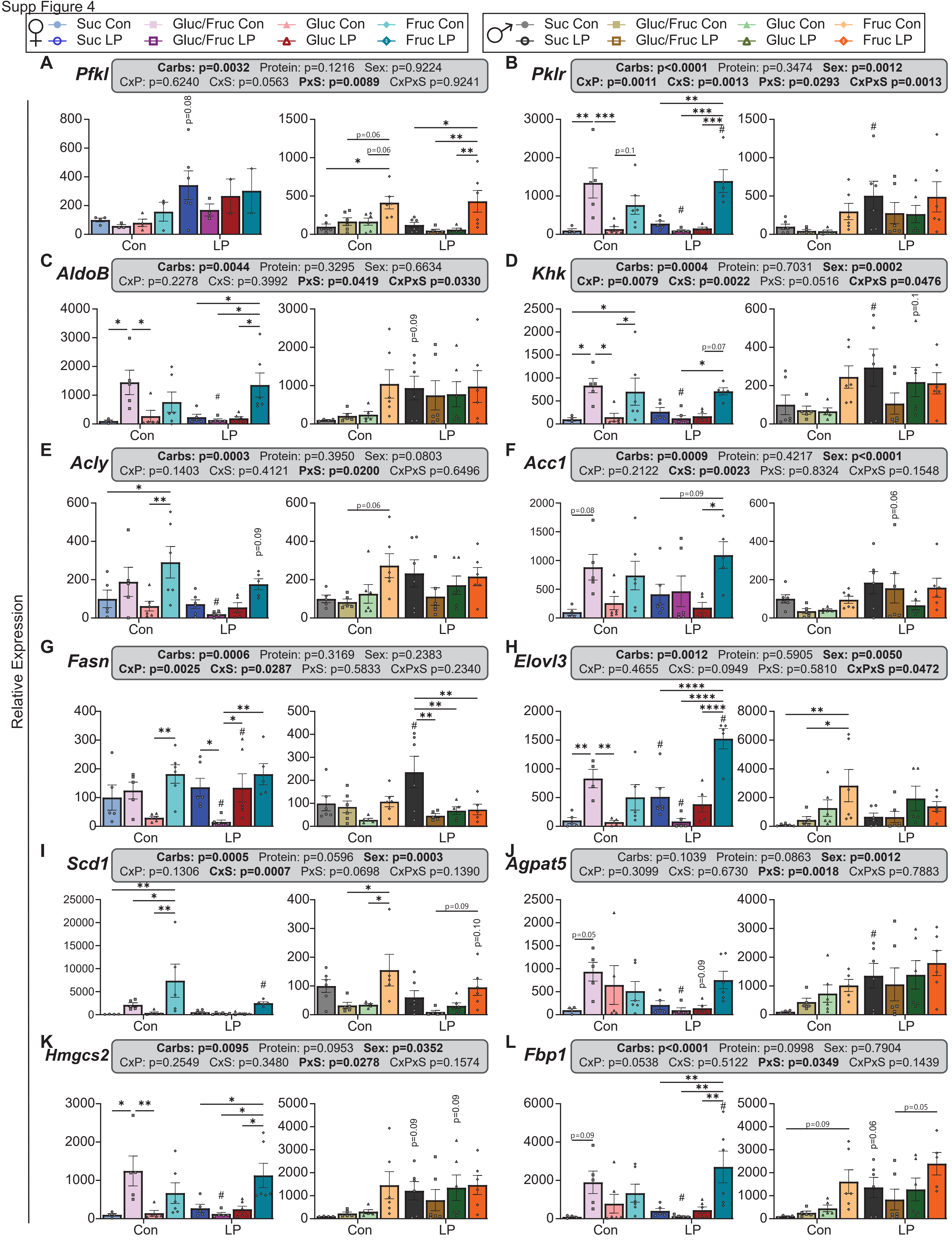

**Figure S4: Expression of liver metabolism genes are increased by low protein and fructose-only diets.**

Individual results for liver qPCRs in the heat map in Fig. 4F; genes measured include:

(A-C) Glycolysis genes *Pfkl* (A), *Pklr* (B), and *AldoB* (C)

(C-D) Fructolysis genes *AldoB* (C) and *Khk*

(E-J) *De novo* lipogenesis genes *Acly* (E), *Acc1* (F), *Fasn* (G), *Elovl3* (H), *Scd1* (I), *Agpat5* (J)

(K) Ketogenic gene *Hmgcs2*

(L) Gluconeogenic gene *Fbp1*

All comparisons in this figure: asterisks denote differences between carbohydrate types within control or LP diets with *$p<0.05$, **$p<0.01$, ***$p<0.001$, ****$p<0.0001$, Tukey test post 2-way ANOVA. Hashes denote significant differences ($p<0.05$, Tukey test post 2-way ANOVA) in LP diets compared to their corresponding control counterparts. Exact p-values are provided in Table S3.

A

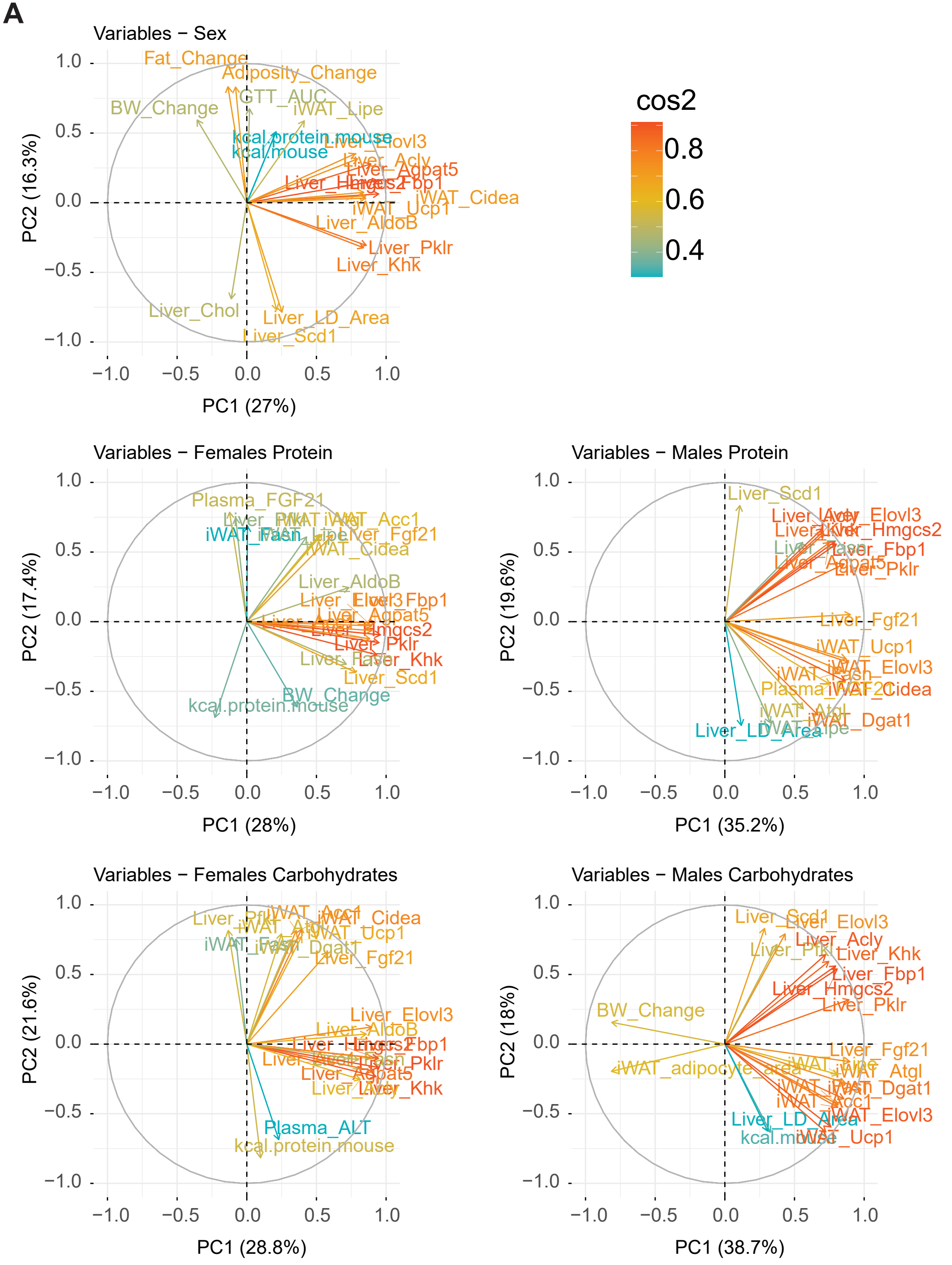
Variables − Sex
PC1 (27%)
PC2 (16.3%)
Fat_Change
Adiposity_Change
GTT_AUC
iWAT_Lipe
BW_Change
kcal.protein.mouse
kcal.mouse
Liver_Elovl3
Liver_Acly
Liver_Agpat5
iWAT_Cidea
iWAT_Ucp1
Liver_AldoB
Liver_Pklr
Liver_Khk
Liver_Chol
Liver_LD_Area
Liver_Scd1
cos2
0.8
0.6
0.4
Variables − Females Protein
PC1 (28%)
PC2 (17.4%)
Plasma_FGF21
iWAT_Acc1
Liver_Fgf21
iWAT_Cidea
Liver_AldoB
Liver_Agpat5
Liver_Hmgcs2
Liver_Pklr
Liver_Khk
Liver_Scd1
BW_Change
kcal.protein.mouse
Variables − Males Protein
PC1 (35.2%)
PC2 (19.6%)
Liver_Scd1
Liver_Elovl3
Liver_Hmgcs2
Liver_Fbp1
Liver_Agpat5
Liver_Pklr
Liver_Fgf21
iWAT_Ucp1
iWAT_Elovl3
iWAT_Cidea
iWAT_Dgat1
Liver_LD_Area
Variables − Females Carbohydrates
PC1 (28.8%)
PC2 (21.6%)
iWAT_Cidea
iWAT_Ucp1
Liver_Fgf21
Liver_Elovl3
Liver_AldoB
Liver_Pklr
Liver_Agpat5
Liver_Khk
Plasma_ALT
kcal.protein.mouse
Variables − Males Carbohydrates
PC1 (38.7%)
PC2 (18%)
Liver_Scd1
Liver_Elovl3
Liver_Acly
Liver_Khk
Liver_Fbp1
Liver_Hmgcs2
Liver_Pklr
BW_Change
Liver_Fgf21
iWAT_adipocyte_area
iWAT_Atgl
iWAT_Dgat1
iWAT_Acc1
iWAT_Elovl3
Liver_LD_Area
iWAT_Ucp1

**Figure S5: Body composition and liver and iWAT gene expression drive differences between groups.**

(A) Principal component analysis was performed, and the contribution of each variable to each dimension calculated. The black box brackets variables that have the strongest contribution to dimensions 1 and 2 within each of the comparisons shown in Figure 5B-D.

Figure S6

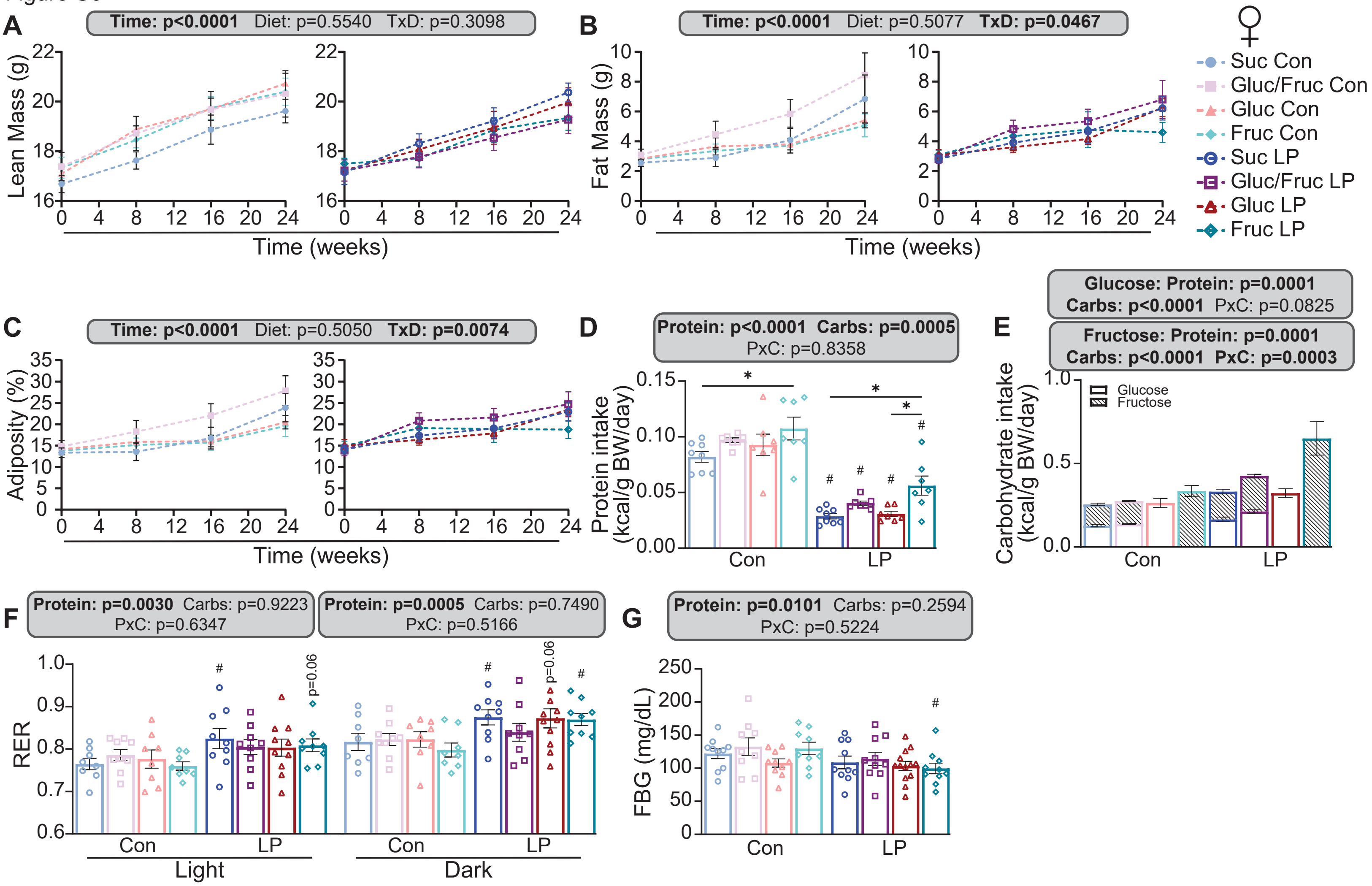

A
Time: p<0.0001 Diet: p=0.5540 TxD: p=0.3098
Lean Mass (g)
Time (weeks)
B
Time: p<0.0001 Diet: p=0.5077 TxD: p=0.0467
Fat Mass (g)
Time (weeks)
Suc Con
Gluc/Fruc Con
Gluc Con
Fruc Con
Suc LP
Gluc/Fruc LP
Gluc LP
Fruc LP
C
Time: p<0.0001 Diet: p=0.5050 TxD: p=0.0074
Adiposity (%)
Time (weeks)
D
Protein: p<0.0001 Carbs: p=0.0005
PxC: p=0.8358
Protein intake (kcal/g BW/day)
Con
LP
E
Glucose: Protein: p=0.0001
Carbs: p<0.0001 PxC: p=0.0825
Fructose: Protein: p=0.0001
Carbs: p<0.0001 PxC: p=0.0003
Carbohydrate intake (kcal/g BW/day)
Glucose
Fructose
Con
LP
F
Protein: p=0.0030 Carbs: p=0.9223
PxC: p=0.6347
Protein: p=0.0005 Carbs: p=0.7490
PxC: p=0.5166
RER
p=0.06
Con
LP
Light
Dark
G
Protein: p=0.0101 Carbs: p=0.2594
PxC: p=0.5224
FBG (mg/dL)
Con
LP

**Figure S6: Fat mass and adiposity accretion is worsened in Gluc/Fruc-fed APP/PS1 mice.**

(A) Lean mass over the course of the experiment in female APP/PS1 mice.

(B) Fat mass over the course of the experiment in female APP/PS1 mice.

(C) Adiposity over the course of the experiment in female APP/PS1 mice.

(D) Protein intake normalized to body weight, calculated from home cage food consumption.

(E) Glucose (solid) and fructose (striped) consumption normalized to body weight, calculated from home cage food consumption.

All comparisons in this figure: asterisks denote differences between carbohydrate types within control or LP diets with *$p<0.05$, **$p<0.01$, ***$p<0.001$, ****$p<0.0001$, Tukey test post 2-way ANOVA. Hashes denote significant differences ($p<0.05$, Tukey test post 2-way ANOVA) in LP diets compared to their corresponding control counterparts. Exact p-values are provided in Table S3.

Figure S7

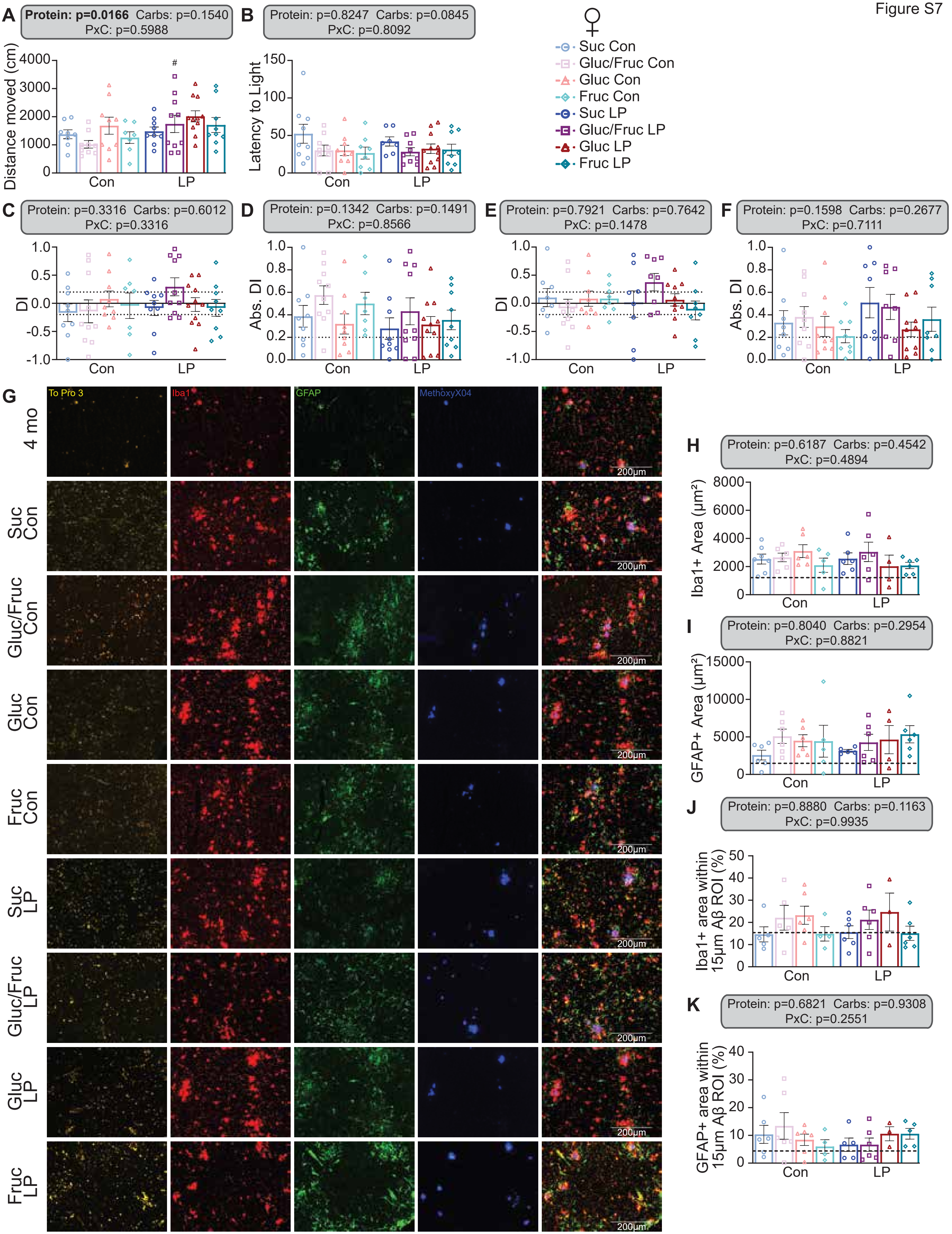

A
Protein: p=0.0166 Carbs: p=0.1540
PxC: p=0.5988
Distance moved (cm)
#
Con
LP
B
Protein: p=0.8247 Carbs: p=0.0845
PxC: p=0.8092
Latency to Light
Con
LP
♀
Suc Con
Gluc/Fruc Con
Gluc Con
Fruc Con
Suc LP
Gluc/Fruc LP
Gluc LP
Fruc LP
C
Protein: p=0.3316 Carbs: p=0.6012
PxC: p=0.3316
DI
D
Protein: p=0.1342 Carbs: p=0.1491
PxC: p=0.8566
Abs. DI
E
Protein: p=0.7921 Carbs: p=0.7642
PxC: p=0.1478
DI
F
Protein: p=0.1598 Carbs: p=0.2677
PxC: p=0.7111
Abs. DI
G
To Pro 3
Iba1
GFAP
MethoxyX04
4 mo
Suc Con
Gluc/Fruc Con
Gluc Con
Fruc Con
Suc LP
Gluc/Fruc LP
Gluc LP
Fruc LP
200μm
H
Protein: p=0.6187 Carbs: p=0.4542
PxC: p=0.4894
Iba1+ Area (μm²)
I
Protein: p=0.8040 Carbs: p=0.2954
PxC: p=0.8821
GFAP+ Area (μm²)
J
Protein: p=0.8880 Carbs: p=0.1163
PxC: p=0.9935
Iba1+ area within 15μm Aβ ROI (%)
K
Protein: p=0.6821 Carbs: p=0.9308
PxC: p=0.2551
GFAP+ area within 15μm Aβ ROI (%)

**Figure S7: Gliosis, anxiety, and object recognition are not impacted by carbohydrate type or protein level.**

(A-B) Open field test (A) and dark/light box (B) was performed to assess anxiety, and distance moved and latency to light quantified, respectively.

(C-F) Novel object recognition was performed. 1 hour after familiar object training, a short-term memory test was performed and discrimination index (DI; C) and absolute discrimination index (Abs. DI; D) quantified. 24 hours later, long-term memory test was performed (E-F).

(G-K) Co-staining between Iba1 (red), GFAP (green), and amyloid (blue) was performed with To Pro 3 used as a nuclear stain and imaged at 20x on an Evos Autofl microscope. Representative images are shown in (G). Total Iba1+ area (H) and GFAP+ area (I) within the field of view was quantified. A 15μm region of interest (ROI) was drawn around each plaque, and the percent of that ROI that was Iba1+ (J) or GFAP+ (K) was measured. Dotted lines indicate levels found in 4-month-old APP/PS1 mice.

n=5-9. All comparisons in this figure: asterisks denote differences between carbohydrate types within control or LP diets with *$p<0.05$, **$p<0.01$, ***$p<0.001$, ****$p<0.0001$, Tukey test post 2-way ANOVA. Hashes denote significant differences ($p<0.05$, Tukey test post 2-way ANOVA) in LP diets compared to their corresponding control counterparts. Exact p-values are provided in Table S3.

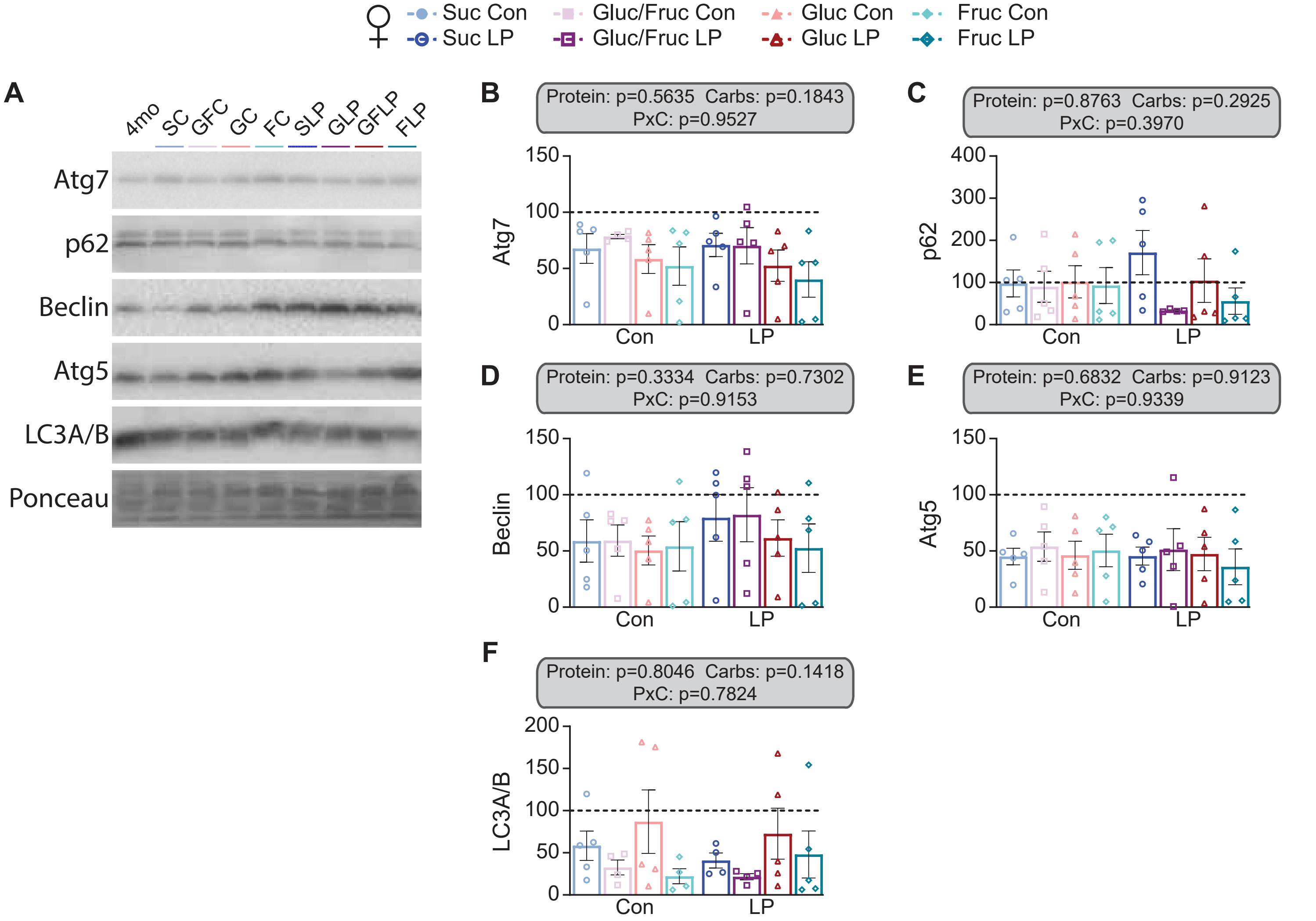
Suc Con
Suc LP
Gluc/Fruc Con
Gluc/Fruc LP
Gluc Con
Gluc LP
Fruc Con
Fruc LP
A
4mo
SC
GFC
GC
FC
SLP
GLP
GFLP
FLP
Atg7
p62
Beclin
Atg5
LC3A/B
Ponceau
B
Protein: p=0.5635 Carbs: p=0.1843
PxC: p=0.9527
Atg7
Con
LP
C
Protein: p=0.8763 Carbs: p=0.2925
PxC: p=0.3970
p62
Con
LP
D
Protein: p=0.3334 Carbs: p=0.7302
PxC: p=0.9153
Beclin
Con
LP
E
Protein: p=0.6832 Carbs: p=0.9123
PxC: p=0.9339
Atg5
Con
LP
F
Protein: p=0.8046 Carbs: p=0.1418
PxC: p=0.7824
LC3A/B
Con
LP

**Figure S8: Autophagy is not altered by carbohydrate type or protein level.**

A) Representative Western blot images.

B-F) Quantification of Western blots for autophagy proteins Atg7 (B), p62 (B), Beclin (C), Atg5 (D), and LC3A/B (E). Dotted lines indicate levels found in 4-month-old APP/PS1 mice.

N=4-5. Exact p-values are provided in Table S3.

**Table Legends**

**Table S1:** Composition and calorie content for diets used in this study.

**Table S2:** Primer sequences used for RT-qPCR.

**Table S3:** Antibodies used for Western blotting and immunohistochemistry.

**Table S4:** P-values from two-way ANOVAs and corresponding multiple comparisons.

**Table S5:** Z-score metrics
